# CRISPR-Cas9-assisted native end-joining editing offers a simple strategy for efficient genetic engineering in *Escherichia coli*

**DOI:** 10.1101/605246

**Authors:** Chaoyong Huang, Tingting Ding, Jingge Wang, Xueqin Wang, Jialei Wang, Lin Zhu, Changhao Bi, Xueli Zhang, Xiaoyan Ma, Yi-Xin Huo

**Affiliations:** Key Laboratory of Molecular Medicine and Biotherapy, School of Life Sciences, Beijing Institute of Technology, No. 5 South Zhongguancun Street, Beijing, PR China, 100081; Tianjin Institute of Industrial Biotechnology, Chinese Academy of Sciences, Tianjin, PR China, 300308; UCLA Institute of Advancement (Suzhou), 10 Yueliangwan Road, Suzhou Industrial Park, Suzhou, PR China, 215123

**Keywords:** *Escherichia coli*, genetic engineering, CRISPR/Cas9, end-joining, Large fragment deletion

## Abstract

Unlike eukaryotes, bacteria are less proficient in homologous recombination (HR) and non-homologous end joining (NHEJ). All existing genomic editing methods for *Escherichia coli* rely on exogenous HR or NHEJ systems to repair DNA double-strand breaks (DSBs). Although an *E. coli* native end-joining (ENEJ) system has been reported, its potential in chromosomal engineering has not yet been explored. Here, we present a CRISPR-Cas9-assisted native end-joining editing and show that ENEJ-dependent DNA repair can be used to conduct rapid and efficient knocking-out of *E. coli* genomic sequence of up to 83 kb. Moreover, the positive rate and editing efficiency is independent of high-efficiency competent cells. The method requires neither exogenous DNA repair systems nor introduced editing template. The Cas9 complex is the only foreign element in this method. This study is the first successful engineering effort to utilize ENEJ mechanism in genomic editing and provides an effective strategy for genetic engineering in bacteria that are inefficient in HR and NHEJ.

**Significance:** The application in prokaryotes is difficult because of the weak homologous recombination and non-homologous end joining systems. *E. coli*, as the most-used prokaryote in metabolic engineering, has no NHEJ system. All existing genomic editing methods for *E. coli* rely on exogenous HR or NHEJ systems to repair double-strand breaks introduced by CRISPR/Cas9. In this report, we firstly demonstrate that the weak and previously ignored end-joining mechanism in *E. coli* can be used for efficient large-scale genetic engineering assisted by CRISPR/Cas9. Our efforts greatly simplify the genomic editing procedure of *E. coli* and provide an effective strategy for genetic engineering in bacteria that are inefficient in HR and NHEJ.

## Introduction

The ability to easily and efficiently edit the bacterial genome is highly desirable in metabolic engineering to produce valuable chemicals (1–4). Creating double strand breaks (DSBs) is a key step in driving genomic engineering (5). Novel applications of clustered regularly interspaced short palindromic repeats (CRISPR) and CRISPR-associated protein 9 (Cas9) can generate DSBs at desired genomic loci (6,7). Under the guidance of a trans-activating crRNA (tracrRNA):crRNA duplex or an engineered single-guide RNA (sgRNA), the Cas9 protein recognizes and cleaves a target DNA sequence with a required protospacer adjacent motif (PAM) (6,8). To survive, it is necessary for bacteria to repair the DSBs via homologous recombination (HR) or non-homologous end joining (NHEJ). Genomic editing events occur during DSBs repair.

NHEJ is the prevalent DSB repair mechanism. It joins unrelated DNA ends and creates new genomic combinations between sequences that do not share homology (9–11). Unlike eukaryotes, bacteria are less proficient in or incapable of NHEJ (12). Some bacteria, including phylogenetically distant *Mycobacteria* and *Bacillus subtilis*, possess a rudimentary NHEJ machinery consisting of two proteins, namely the multifunctional ATP-dependent Ligase-D and Ku protein that collectively protect, process, and ligate DNA ends (13,14). Other bacteria, such as the prevalent gut commensal *Escherichia coli* where Ligase-D-like and Ku-like proteins have not been found are generally accepted to be NHEJ-free (12) and rely only on HR to repair DSBs. *E. coli* is frequently used as a negative control for NHEJ experiments in other bacteria (15).

Generally, the artificial DSBs in *E. coli* must be repaired by RecA-mediated HR with a homologous sequence as the editing template (6). However, the native HR pathway is commonly recognized to be of low efficiency in repairing CRISPR-Cas9-introduced DSBs. The introduction of bacteriophage-derived λ-Red recombinases made a breakthrough in the efficiency of making desired mutations in the genome (6,16,17). Recently, *Su et al*. and *Zheng et al*. introduced a *Mycobacteria*-derived NHEJ pathway into *E. coli* to repair the DSBs generated by CRISPR-Cas9 at desired genomic loci and demonstrated the rapid and efficient inactivation of bacterial genes in a HR-independent manner (1,18). According to the views of *Su et al*. and *Zheng et al*., *E. coli* acquired end-joining activity after overexpressing *Mycobacteria*-derived Ligase-D and Ku protein. Taken together, all existing genomic editing methods of *E. coli* rely on HR pathways, generally assisted by λ-Red recombinases, or exogenous NHEJ pathways to repair DSBs generated by CRISPR-Cas9.

A Ligase-D- and Ku-independent *E. coli* native end-joining (ENEJ) mechanism differs from classic NHEJ mechanism has been reported in *E. coli* and was named as alternative end-joining (A-EJ) (11). In that study, DSBs generated by restriction endonuclease I-SceI were repaired by A-EJ with an efficiency of 10^−5^. However, we have not seen any report on the application of ENEJ in genetic engineering. To fill in the gap and add to the toolbox of genetic engineers, we developed <u>C</u>RISPR-Cas9-assisted <u>n</u>ative <u>e</u>nd-joining <u>e</u>diting (CNEE), in which the ENEJ was utilized to repair DSBs generated by CRISPR-Cas9. Using a systematically optimized protocol, we demonstrated that ENEJ could be used to enable the rapid and efficient deletion of *E. coli* genomic sequence up to 83 kb in length. The method requires neither editing template and λ-Red recombinases nor exogenous NHEJ associated proteins, thus realizing the *E. coli* genomic editing with the lowest number of foreign elements. The method has a similar, if not higher, editing efficiency to previous methods that overexpress exogenous DNA repair proteins. Moreover, the method does not require high-competency host strains, greatly decreasing the difficulty of genomic editing of wild-type strains.

We hypothesized that the strategies used in our method might be useful for other CRISPR-Cas9-mediated genomic editing methods in which DSBs are repaired by RecA-mediated HR, λ-Red-mediated HR or NHEJ. To verify that, we applied the similar strategies in these methods. After modification, the efficiencies of these methods significantly increased.

Taken together, this work is the first to utilize ENEJ in genomic editing in *E. coli* and demonstrates that genomic editing of prokaryotes can be independent of exogenous DNA repair systems.

## Results

### All existing *E. coli* genomic editing methods are dependent on the exogenous DNA repair system

In *E. coli*, CRISPR-Cas9-introduced DSBs lead to cell death unless repaired in a timely manner by DNA repair pathways (19) such as native RecA-mediated HR or A-EJ (11). CRISPR-Cas9-based genomic editing methods achieve genetic modifications by introducing DSBs in the genome and repairing it via DNA repair pathways. In all existing methods, bacteriophage-derived λ-Red system, a HR system, or *Mycobacteria-*derived NHEJ system is introduced into *E. coli* to increase the efficiency of HR or end-joining.

To compare the performances of existing *E. coli* genomic editing methods, we tested all reported methods for *lacZ* inactivation in the wild-type MG1655 [MG1655 (WT)] strain (Table 1). These methods rely on either λ-Red mediated-HR or NHEJ to repair CRISPR-Cas9-induced DSBs. When using HR-dependent methods, a 20-bp-designed DNA sequence was inserted into the open reading frame (ORF) of the *lacZ* gene to generate a frameshift mutation. When using NHEJ-dependent protocols, the stochastic length of the DNA sequence of the ORF was deleted, thus destroying the *lacZ* gene (Fig. 1a). Edited colonies were verified via blue-white screening, colony PCR, and sequencing. The edited colonies were white in the Luria-Bertani (LB) plate containing IPTG and X-gal, while the WT colonies were blue.

**Table 1.** Test results of reported genomic editing methods.

| Foreign elements | DSB repair mechanisms |  |  |  |  |  |  |  |
| --- | --- | --- | --- | --- | --- | --- | --- | --- |
| | λ-Red-mediated HR | | RecA-mediated HR | | NHEJ | | ENEJ | |
| Cas9 protein | + |  | + |  | + |  | + |  |
| Editing template | + |  | + |  | — |  | — |  |
| Red$\gamma\beta\alpha$ | + | | — | | — | | — | |
| Ku, Ligase-D | — |  | — |  | + |  | — |  |
| Tested methods | PR (%) | EE | PR (%) | EE | PR (%) | EE | PR (%) | EE |
| Method 1 (6) | 65 ($\pm 7.2$) | $2.0 (\pm 0.32) \times 10^{-5}$ | 0.71 ($\pm 0.21$) | $1.0 (\pm 0.22) \times 10^{-7}$ | NA | NA | 0 | 0 |
| Method 2 (24) | 86 ($\pm 6.7$) | $1.5 (\pm 0.14) \times 10^{-5}$ | 5.6 ($\pm 0.79$) | $1.3 (\pm 0.18) \times 10^{-7}$ | NA | NA | 2.8 ($\pm 0.99$) | $5.0 (\pm 1.7) \times 10^{-8}$ |
| Method 3 (16) | 95 ($\pm 3.8$) | $2.3 (\pm 0.26) \times 10^{-4}$ | 0.58 ($\pm 0.11$) | $6.0 (\pm 0.48) \times 10^{-8}$ | NA | NA | 0 | 0 |
| Method 4 (20) | 19 ($\pm 6.6$) | $1.4 (\pm 0.12) \times 10^{-5}$ | 2.1 ($\pm 0.34$) | $3.6 (\pm 0.42) \times 10^{-7}$ | NA | NA | 1.5 ($\pm 0.76$) | $2.0 (\pm 0.90) \times 10^{-8}$ |
| Method 5 (25) | 93 ($\pm 5.4$) | $3.9 (\pm 0.38) \times 10^{-5}$ | 0 | 0 | NA | NA | 0 | 0 |
| Method 6 (26) | 100 ($\pm 0$) | $4.6 (\pm 0.44) \times 10^{-6}$ | 0 | 0 | NA | NA | 0 | 0 |
| Method 7 (1) | NA | NA | NA | NA | 51 ($\pm 8.2$) | $2.7 (\pm 0.30) \times 10^{-7}$ | 0 | 0 |
| Method 8 (18) | NA | NA | NA | NA | 90 ($\pm 6.2$) | $9.9 (\pm 1.6) \times 10^{-7}$ | 0 | 0 |

PR: positive rate, EE: editing efficiency, NA: not available. Data are means ± s.d. from three independent experiments.

**Fig. 1.**
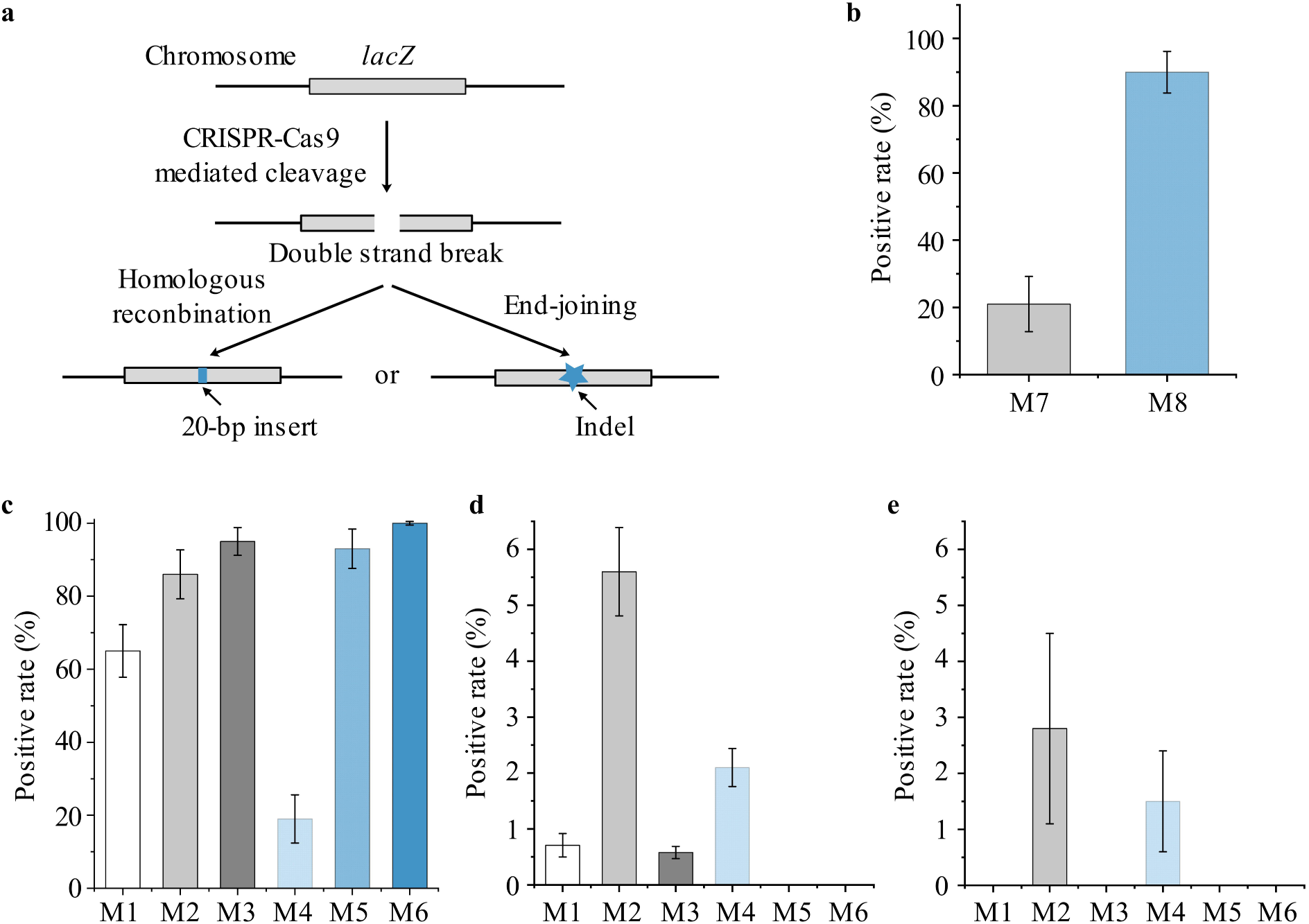
Genomic editing tests of existing methods. (**a**) Schematic of *lacZ* inactivation via HR pathway or end-joining pathway. (**b**) Positive rates of methods 7 and 8 in the presence of overexpressed Ligase-D and Ku protein. (**c**) Positive rates of methods 1-6 in the presence of editing template and overexpressed λ-Red recombinases. (**d**) Positive rates of methods 1-6 without overexpressed λ-Red recombinases but with editing template. (**e**) Positive rates of methods 1-6 in the absence of both editing template and overexpressed λ-Red recombinases. Data are expressed as means ± s.d. from three independent experiments.

Six λ-Red-dependent methods (methods 1-6) were tested (Table 1). The procedures of these six methods are summarized in Fig. S1-S6, respectively. The results showed that five out of six methods achieved a high positive rate of > 65% (Fig. 1c), and the editing efficiency ranged from 4.6 × 10^−6^ to 2.3 × 10^−4^ (Table 1, column 2). These findings indicated that CRISPR-Cas9-induced DSBs were efficiently repaired by the λ-Red mediated HR. Then, the same six methods were tested without overexpressing λ-Red recombinases (Table 1, column 3). In the absence of λ-Red recombinases, RecA was the only recombinase involved in HR that repaired CRISPR-Cas9-induced DSBs. As shown in Fig. 1d, all methods achieved a low positive rate of < 6%, and methods 5 and 6 even showed no positive colonies (white colonies). For methods 1-4, the editing efficiency correspondingly decreased to 6.0 × 10^−8^-3.6 × 10^−7^ (Table 1, column 3). The six methods were also tested in the absence of both λ-Red recombinases and DNA template (Table 1, column 5). The results showed that only methods 2 and 4 generated positive colonies, with rates of 2.8% and 1.5%, respectively (Fig. 1e), and the editing efficiency were 5.0 × 10^−8^ and 2.0 × 10^−8^, respectively (Table 1, column 4).

Two NHEJ-dependent methods (methods 7 and 8) were also tested (Table 1), and the procedures of these two methods are summarized in Fig. S7 and S8, respectively. Both methods achieved a high positive rate of > 50% (Fig. 1b). However, the editing efficiencies of the two methods were only 2.7 × 10^−7^ and 9.9 × 10^−7^, respectively (Table 1, column 4). Methods 7 and 8 were then tested in the absence of overexpressed Ligase-D and Ku proteins. The results showed that no positive colonies (white colonies) were obtained (Table 1, column 5).

### Establishment of CNEE

No previous reports have reported genomic editing in the absence of exogenous DNA repair system and editing template. However, by testing all previous methods for genomic editing, for the first time, we show that ENEJ may be potentially used in genomic editing (Table 1, column 5). To fully utilize ENEJ-mediated DSB repair and increase the efficiency of CRISPR-Cas9, we established CNEE and conducted systematic optimizations (Table S1).

We constructed a two-plasmid system in which the Cas9 and sgRNA were expressed by different plasmids, namely p-P_BAD_-*cas9* and p-P_BAD_-sgRNA-X (X refers to the target DNA) (Fig. 2). The two-plasmid system was more convenient in plasmid construction than the one-plasmid system (16,20). The Cas9 used here was xCas9-3.7, an evolved SpCas9 variant with broad PAM compatibility and high DNA specificity (7). Furthermore, sgRNA other than tracrRNA:crRNA duplex was applied to increase the performance and stability of the CRISPR-Cas9 system (1,8). The combination of xCas9-3.7 and sgRNA increased the efficiency of the CRISPR-Cas9 system, thus reducing the survival rate of WT cells. The expressions of both Cas9 and sgRNA were controlled by an inducible promoter, thus the CRISPR-Cas9 system functioned only when an inducer was added. The L-arabinose-induced P_BAD_ promoter and two IPTG-induced promoters, namely P_T5_ and P_L_lacO_1_, were tested individually. We found that the P_BAD_ promoter was stricter than both P_T5_ and P_L_lacO_1_, so we used the P_BAD_ promoter for Cas9 and sgRNA expressions (Table S1). Then, we optimized the working concentration of L-arabinose. Different concentrations of L-arabinose, ranging from 1-30 mM, were tested to induce the CRISPR-Cas9 system, and L-arabinose at a concentration of 20 mM led to the highest positive rate and editing efficiency (Table S1). We found that L-arabinose at low concentrations (*i.e*., 1 mM) were inefficient to induce the CRISPR-Cas9 system; and that high-concentration L-arabinose (*i.e*., 30 mM) resulted in higher rate of cell death, which might be due to the toxicity effect of high amounts of the Cas9 protein (21). Culture temperature was also an important factor in genomic editing. Low culture temperatures resulted in a reduction in cell growth rate, whereas high culture temperatures reduced the stability of plasmid p-P_BAD_-sgRNA-X, which is temperature sensitive (22). Different culture temperatures, ranging from 25°C to 37°C, were tested for genomic editing, and 30°C showed the highest positive rate and editing efficiency (Table S1). In addition, we also optimized the culture time and medium in order to further improve the positive rate and editing efficiency (Table S1).

**Fig. 2.**
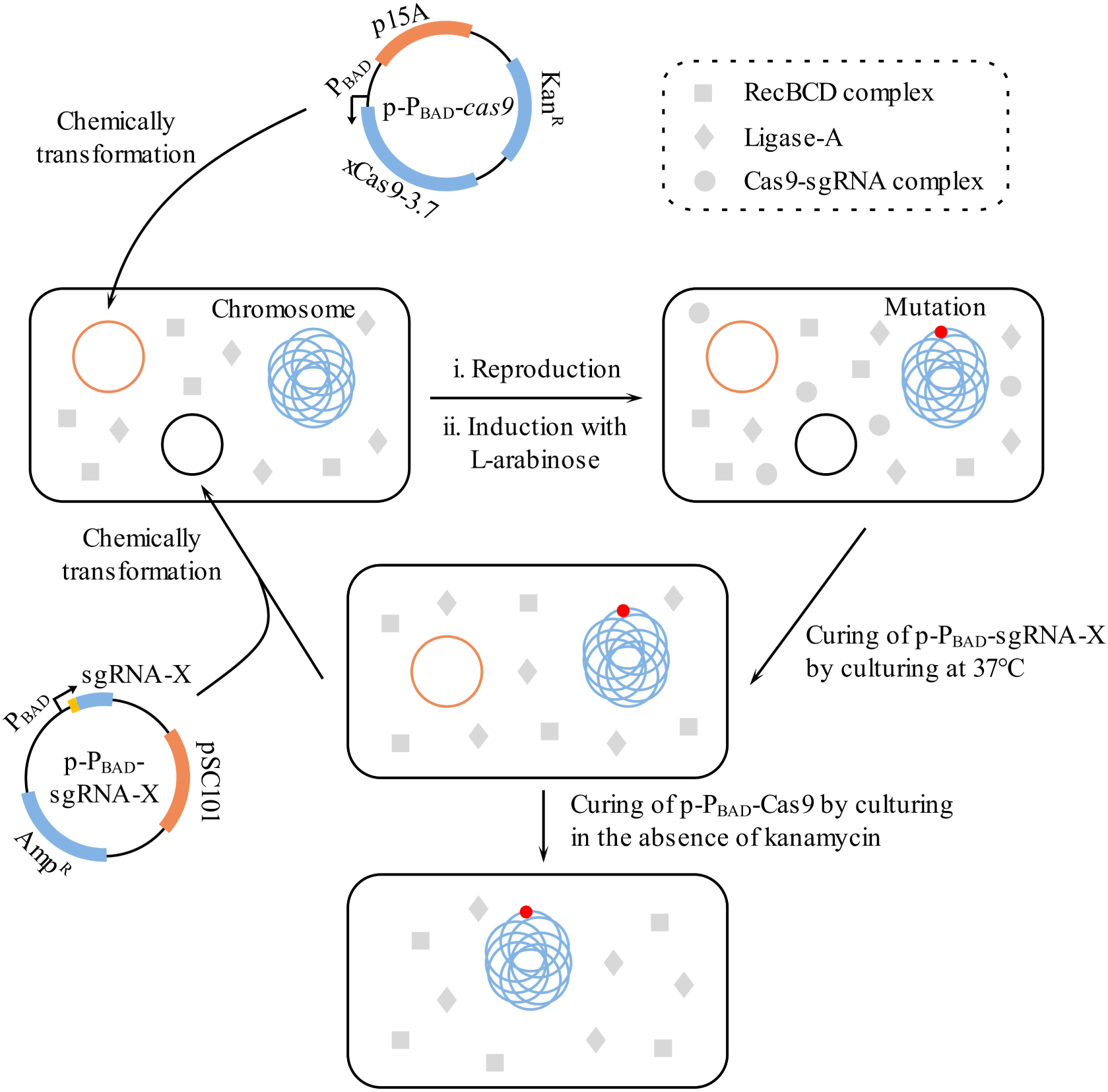
Schematic of CNEE. Step 1: Plasmids p-P_BAD_-*cas9* and p-P_BAD_-sgRNA-X were transformed into a host strain. Step 2: a transformant was cultured for reproduction before inducing genomic editing by adding L-arabinose. Step 3: Plasmids in the edited strain were removed to obtain a plasmid-free strain.

The schematic procedures of the optimized CNEE are presented in Fig. 2. Detailed procedures are provided in Fig. 9. First, the plasmid p-P_BAD_-*cas9* expressing Cas9 and the plasmid p-P_BAD_-sgRNA expressing sgRNA were transformed into the target strain. Then, after one of the transformants was cultured for a period of time, L-arabinose was applied to induce the expression of Cas9 and sgRNA, resulting in site-specific generations of DSBs and deletion mutations. Briefly, the Cas9-sgRNA complex binds and cleaves the target DNA strands and generates DSBs at desired genomic loci. The ENEJ pathway repairs the DSBs, and the cells survive the CRISPR-Cas9 mediated cleavage but carry stochastic deletion in the target genomic loci (11). Finally, a small fraction of the culture was plated in a selection plate, and the edited strains were verified by colony PCR and sequencing. The plasmid p-P_BAD_-sgRNA-X was removed by increasing the cultivation temperature to 37-42°C (Fig. S10a), and a new p-P_BAD_-sgRNA-X plasmid targeting a different genomic locus was introduced for the next round of genomic editing. Finally, the plasmid p-P_BAD_-Cas9 was cured by culturing in LB antibiotic-free medium (Fig. S10b).

### CNEE is convenient for gene inactivation

In principle, the ENEJ system repairs CRISPR-Cas9-induced DSBs and results in stochastic genomic sequence deletion, thus disrupting the target ORF (Fig. 3a). We first inactivated the *lacZ* gene in the MG1655 (WT) strain (Fig. 3b). The plasmid p-P_BAD_-sgRNA-*lacZ* containing the spacer that target the *lacZ* gene was transformed into MG1655 (WT) harboring plasmid p-P_BAD_-*cas9*; the mutations of the *lacZ* gene in the resulting strains was examined by blue-white screening, colony PCR, and sequencing.

**Fig. 3.**
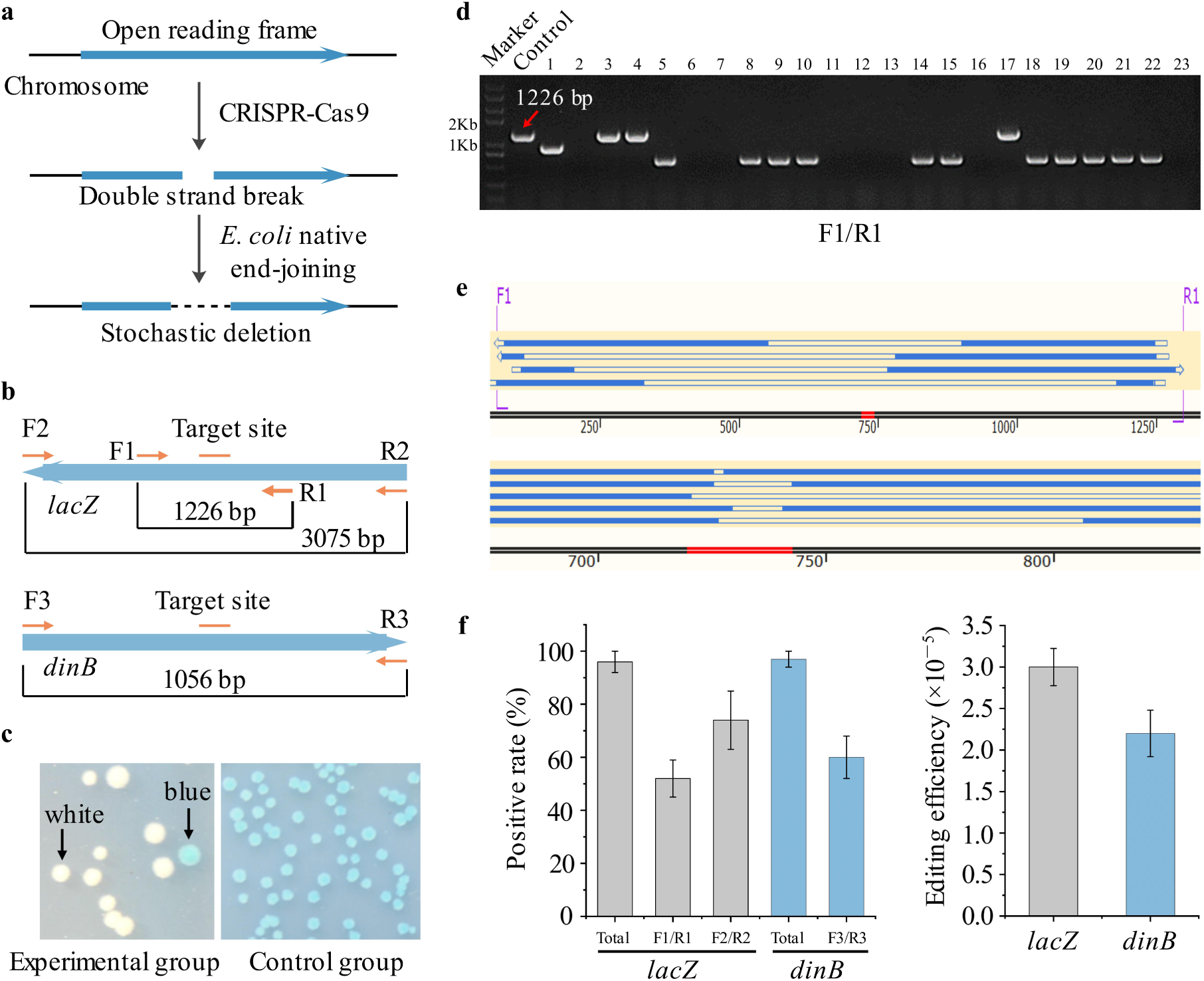
CNEE based gene inactivation. (**a**) Principle of gene inactivation. (**b**) Genes that were inactivated. (**c**) Representative results of blue-white screening of *lacZ* inactivation experiment. (**d**) Representative PCR results of *lacZ* inactivation experiment. White colonies were randomly picked to conduct PCR screening with a blue colony served as the control. F1/R1 were used as primers. (**e**) Representative sequencing results of *lacZ* inactivation experiment. (**f**) Positive rates and editing efficiencies of *lacZ* inactivation and *dinB* inactivation. “Total” represents all edited colonies. “Fn/Rn (n=1, 2, 3)” represent colonies that contain sequence deletion within the corresponding primer pairs. Data are expressed as means ± s.d. from three independent experiments.

Almost all colonies in the experimental group turned white, indicating that most cells surviving CRISPR-Cas9-mediated cleavage were edited, whereas all colonies in the control group were blue in color, indicating that genomic editing did not occur in the absence of CRISPR-Cas9 (Fig. 3c). The positive rate was 96% and the editing efficiency was 3 × 10^−5^ (Fig. 3f). One hundred white colonies were randomly picked for PCR screening using primer pairs F1/R1 and F2/R2. Theoretically, an agarose gel electrophoresis band could be obtained if the sequence deletion was within the complementary regions of primer pair. The results showed that using primer pair F1/R1, 52% of the white colonies generated one band, indicating that 52% of total positive colonies carried a sequence deletion within the complementary regions of F1/R1 (Fig. 3f). Part of the PCR results are shown in Fig. 3d. The control group showed a band of 1,226 bp in size, the experimental group, *i.e*. Nos. 1 and 5, exhibited a band that are obviously less than 1,226 bp in size. For colonies that did not exhibit a band, *i.e*. Nos. 2 and 6, deletions beyond the complementary regions of F1/R1 were observed. Part of the sequencing results are shown in Fig. 3e. Deletions of different lengths that were within the complementary regions of F1/R1 were observed. Similarly, using primer pair F2/R2, 74% of the white colonies exhibited one band of less than 3,075 bp, indicating that 74% of total positive colonies carried a sequence deletion within the ORF of *lacZ* gene (Fig. 3f). For colonies that did not exhibit a band, deletions beyond the ORF of *lacZ* gene were observed. Generally, these colonies were excluded because sequence deletions that exceed the length of the ORF might impair the function of neighbouring genes. We still found blue colonies in the experimental group, so we examined 10 of these by colony PCR and sequencing. All tested blue colonies were determined to have an intact *lacZ* gene, generally, these colonies are called ‘escapers’. Although ‘escapers’ still existed, these only accounted for 4% of all the colonies, making colony screening much less tedious.

Using the same approach, we then iteratively knocked out *dinB*, a gene with an ORF that was smaller than *lacZ* (Fig. 3b). One hundred colonies in the experimental group were randomly picked for PCR screening and sequencing, with F3/R3 as primers. The positive rate was 97%, and the editing efficiency was 2.2 × 10^−5^. All colonies that were not positive (3%) were proven to have an intact *dinB* gene. Among the positive colonies, 60% carried a sequence deletion within the ORF of *dinB* (Fig. 3f).

### CNEE is efficient in knocking out large fragments

Then, CNEE was applied for large-fragment knocking-out in MG1655 (WT), that is, an 81-kb large fragment (3,609,883-3,690,267) and an 83-kb large fragment (2,348,515-2,431,673). To delete the 81-kb large fragment from the bacterial chromosome, we designed the p-P_BAD_-sgRNA-81kb plasmid which contains two sgRNA expression chimeras (Fig. 4a). The plasmid was transformed into MG1655 (WT) that harbored plasmid p-P_BAD_-*cas9*. Theoretically, under the guidance of two sgRNAs, the Cas9 protein cuts off the fragment from the genome by simultaneously introducing a pair of DSBs in the genome, followed by the re-joining of the distant ends (Fig. 4b). There is no essential gene in the 81-kb fragment, thus the knocking-out of this fragment does not impair the growth of the cell. Flanking the fragment are two essential genes *yhhQ* and *bcsB*, and the deletion of either gene is lethal to the cells (23). Therefore, the survived cells should have intact *yhhQ* and *bcsB*.

**Fig. 4.**
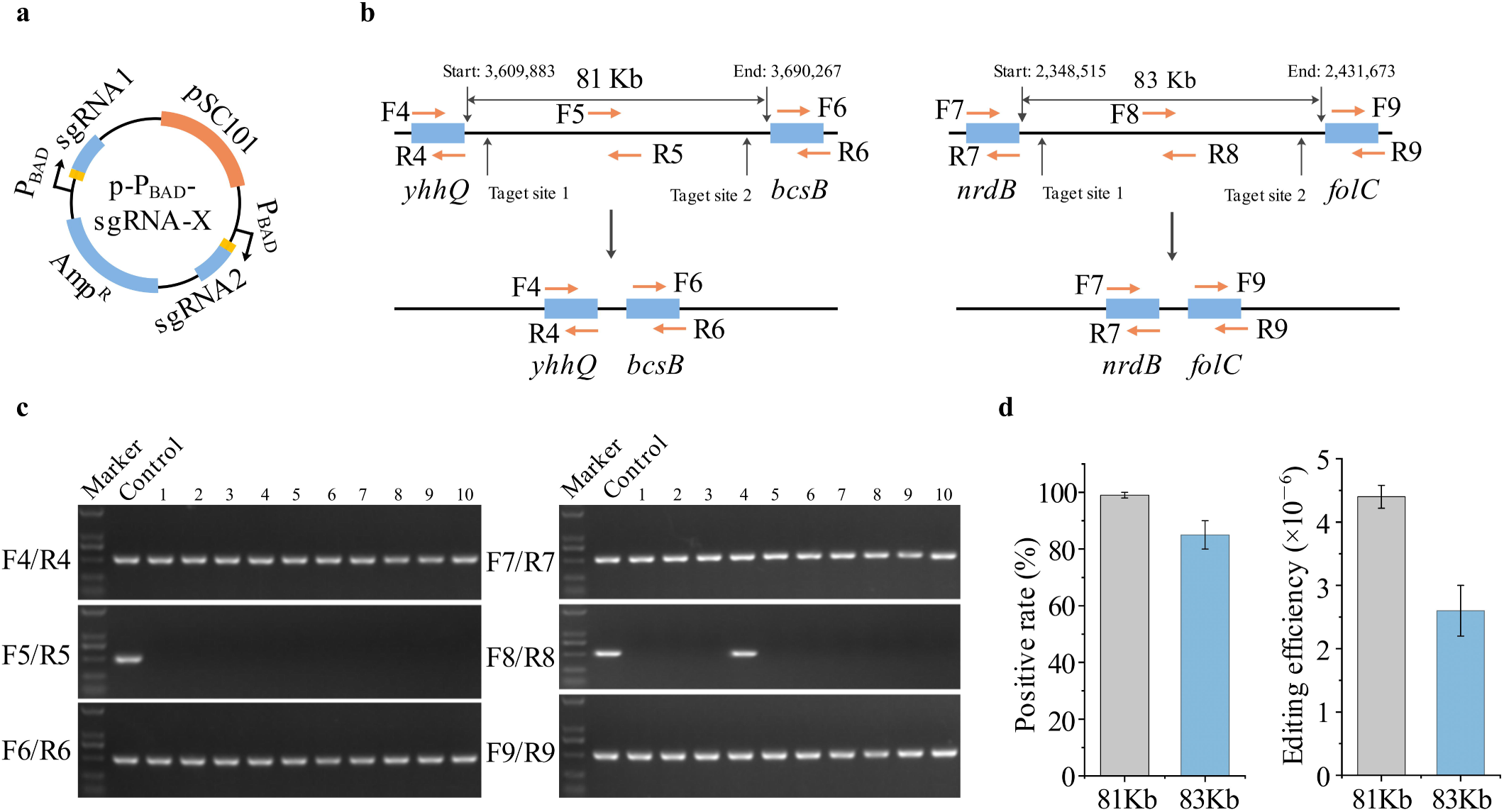
CNEE based large-fragment knocking-out. (**a**) Structure of plasmid p-P_BAD_-sgRNA-X used for large-fragment knocking-out. The plasmid contains two sgRNA expression chimeras. (**b**) Knocking-out of an 81 kb fragment and an 83 kb fragment. (**c**) Representative PCR results of large-fragment knocking-out experiments. Colonies were randomly picked for PCR screening, and a WT colony served as control. (**d**) Positive rates and editing efficiencies of large-fragment knocking-out. Data are expressed as means ± s.d. from three independent experiments.

To check the mutants by colony PCR, three primer pairs were designed to target the regions within the *yhhQ* (F4/R4), the 81-kb fragment (F5/R5), and the *bcsB* (F6/R6) (Fig. 4b). One hundred colonies were randomly selected for PCR screening, and part of agarose gel electrophoresis results are shown in Fig. 4c. The desired mutants had only the first and the third bands, whereas the control [MG1655 (WT)] had all three bands. The positive rate was 99%, and the editing efficiency was 4.4 × 10^−6^ (Fig. 4d).

Using the same approach, we then iteratively knocked out an 83-kb large fragment (Fig. 4b). There is no essential gene in the 83-kb fragment, thus the knocking-out of this fragment does not impair the growth of the cells. Franking the fragment are two essential genes *nrdB* and *folC*, and the deletion of each gene is lethal to the cells (23). Therefore, the survived cells should have intact *nrdB* and *folC*. Primer pairs F7/R7, F8/R8 and F9/R9 were designed to check the positive mutants, and part of agarose gel electrophoresis results are shown in Fig. 4c. The positive rate was 85%, and the editing efficiency was 2.6 × 10^−6^ (Fig. 4d). We also obtained 15% of false positive colonies, *i.e*. the No. 4 in the experimental group (Fig. 4c). Further investigation showed that these false-positive colonies did not lose the 83-kb fragment, but contained stochastic length of deletion in the region of both target sites. The results indicated that the fragment being cut off could, with a certain probability, be re-integrated back into the chromosome through ENEJ.

### CNEE does not require high-efficiency competent cells

Generally, CRISPR-Cas9-based genomic editing methods depend on high-efficiency competent cells to achieve relatively high editing efficiency, and electroporation is commonly used to obtain high transformation efficiency (1,6,16,18,20,24–26). However, highly competent cells, especially chemically competent cells, are not always easy to prepare. Some strains, such as the MG1655 (WT) strain are very difficult to prepare highly competent cells due to their genetic properties (27,28). The CNEE allows transformants to reproduce before inducing genome editing by adding L-arabinose, and thus does not depend on high-competency host strains. Theoretically, as long as one transformant is obtained, genome editing can proceed normally (Fig. 6a). To prove this advantage, we tested the CNEE for gene inactivation using MG1655 (WT) competent cells of different transformation efficiencies. The *lacZ* gene was selected as the target gene. The CaCl_2_ method, Inoue method (27), and electroporation method were used to obtain competent cells of different competencies, ranging from 10^4^ (cfu/μg pUC19) to 10^8^ (cfu/μg pUC19). As expected, there was no apparent difference in either the positive rate or the editing efficiency (Table 2).

**Table 2.** Test results of CNEE using cells with different competencies.

| Competency (cfu/μg pUC19) | PR (%) | EE | Deletion size less than 1 kb (%) |
| --- | --- | --- | --- |
| $10^4$ (CaCl<sub>2</sub> method) | 98 ± 1.4 | $3.2 (\pm 0.21) \times 10^{-5}$ | 55 ± 6.5 |
| $10^5$ (CaCl<sub>2</sub> method) | 96 ± 3.8 | $3.0 (\pm 0.23) \times 10^{-5}$ | 52 ± 7.6 |
| $10^6$ (Inoue method) | 97 ± 2.1 | $3.5 (\pm 0.26) \times 10^{-5}$ | 54 ± 7.1 |
| $10^7$ (Inoue method) | 95 ± 4.2 | $3.3 (\pm 0.32) \times 10^{-5}$ | 53 ± 4.3 |
| $10^8$ (Electroporation method) | 96 ± 3.4 | $3.5 (\pm 0.28) \times 10^{-5}$ | 58 ± 5.7 |

PR: positive rate, EE: editing efficiency. Data are expressed as means ± s.d. from three independent experiments.

In previous methods, the minimal transformation efficiency of competent cells was 10^8^-10^9^ (cfu/μg pUC19) to obtain a relatively high editing efficiency (Table 3). However, in CNEE, a competency of 10^4^ (cfu/μg pUC19) is sufficient to obtain a comparable editing efficiency (Tables 2 and 3).

**Table 3.**
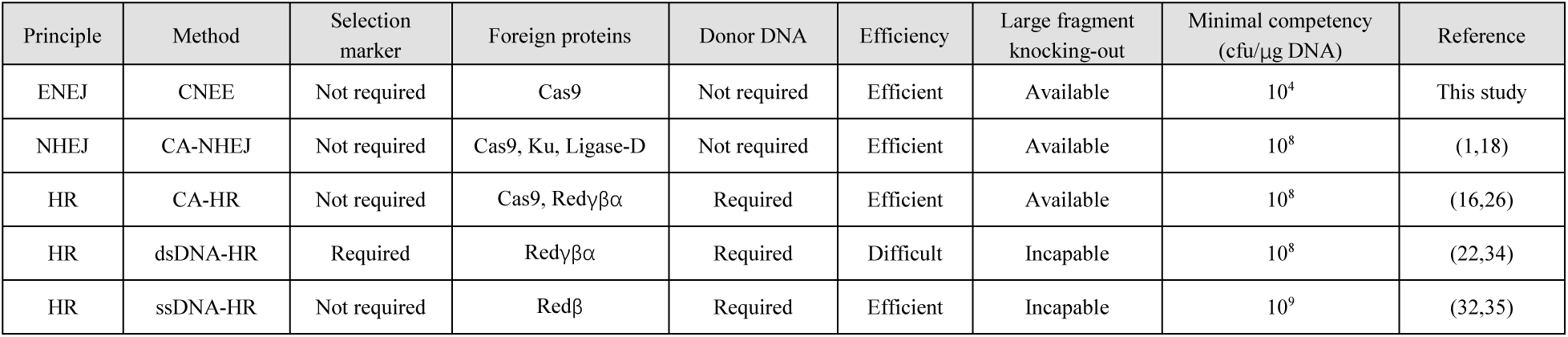
Comparison of CNEE with existing genomic editing methods.

| Principle | Method | Selection marker | Foreign proteins | Donor DNA | Efficiency | Large fragment knocking-out | Minimal competency (cfu/μg DNA) | Reference |
| --- | --- | --- | --- | --- | --- | --- | --- | --- |
| ENEJ | CNEE | Not required | Cas9 | Not required | Efficient | Available | $10^4$ | This study |
| NHEJ | CA-NHEJ | Not required | Cas9, Ku, Ligase-D | Not required | Efficient | Available | $10^8$ | (1,18) |
| HR | CA-HR | Not required | Cas9, Red$\gamma\beta\alpha$ | Required | Efficient | Available | $10^8$ | (16,26) |
| HR | dsDNA-HR | Required | Red$\gamma\beta\alpha$ | Required | Difficult | Incapable | $10^8$ | (22,34) |
| HR | ssDNA-HR | Not required | Redβ | Required | Efficient | Incapable | $10^9$ | (32,35) |

### The DSB repair in CNEE is independent of RecA and LigB, but is dependent on RecBCD

The CNEE system has been proved to be convenient and efficient for gene inactivation and large-fragment knocking-out. However, the detailed mechanism of ENEJ involved in CNEE was unknown. Chayot *et al*. proposed a model for the mechanism of A-EJ, which is the first ENEJ pathway found in *E. coli* (Fig. 5a) (11). In that model, A-EJ relies on the degradation of DNA ends by the RecBCD complex and the ligation by the LigA. In addition, the regions of microhomology (1-9 nt) are essential to promote the ligation of two DNA ends.

**Fig. 5.**
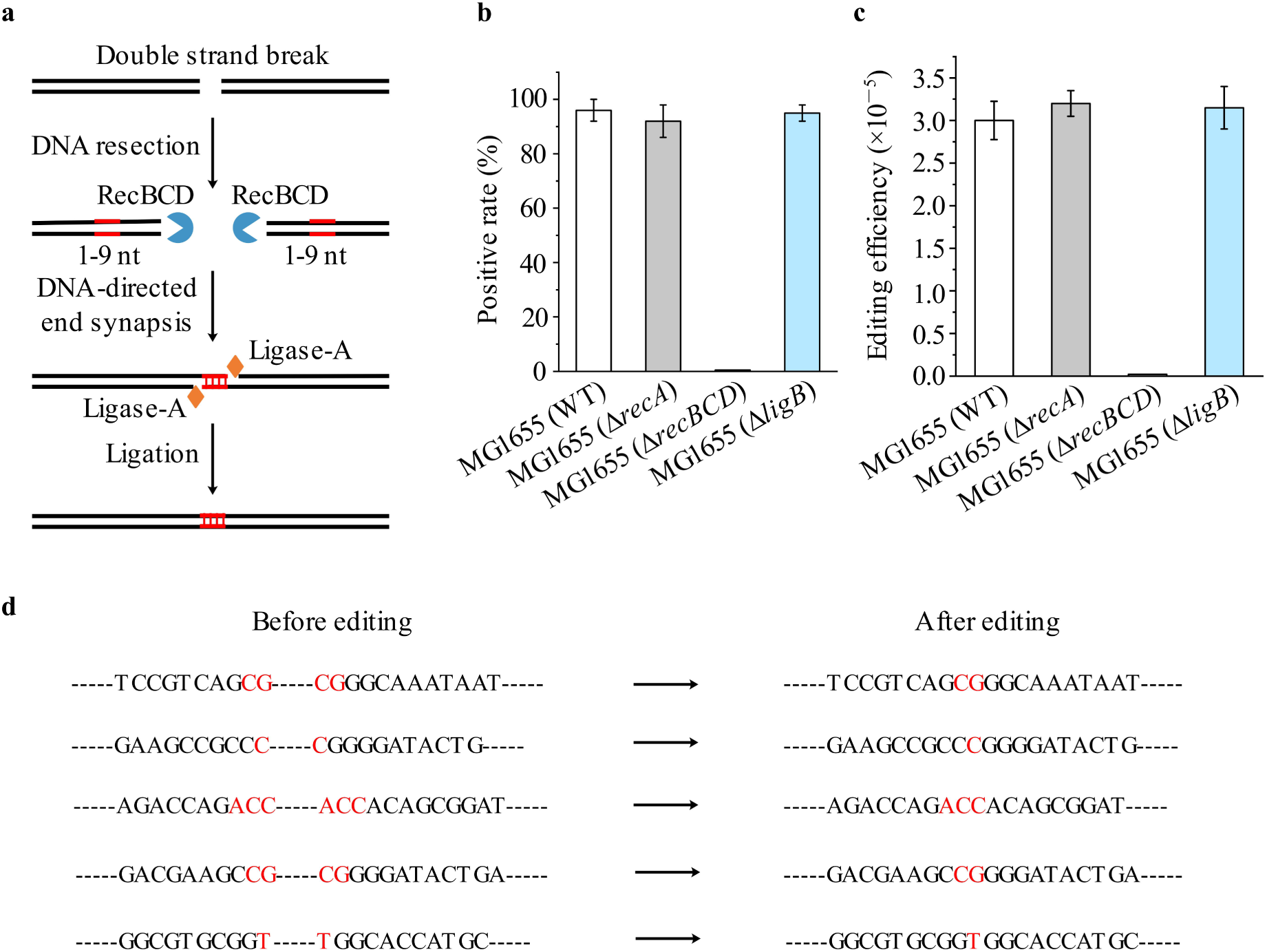
Mechanism of ENEJ involved in CNEE. (**a**) Model of A-EJ mechanism proposed by Chayot *et al*. Step 1: the RecBCD complex binds and degrades DNA ends and generates micro sticky ends in the microhomology regions. Step 2: the sticky ends mediates the synapsis of DNA. Step 3: the Ligase-A repairs the gaps and generates complete dsDNA. (**b**, **c**) Results of *lacZ* inactivation in three strains. *ΔrecA* represents *recA* deficient. *ΔrecBCD* represents *recBCD* deficient. Strains MG1655 (*ΔrecA*) and MG1655 (*ΔrecBCD*) were obtained by CNEE. (**d**) Representative DSB repair events by CNEE. Data are expressed as means ± s.d. from three independent experiments.

**Fig. 6.**
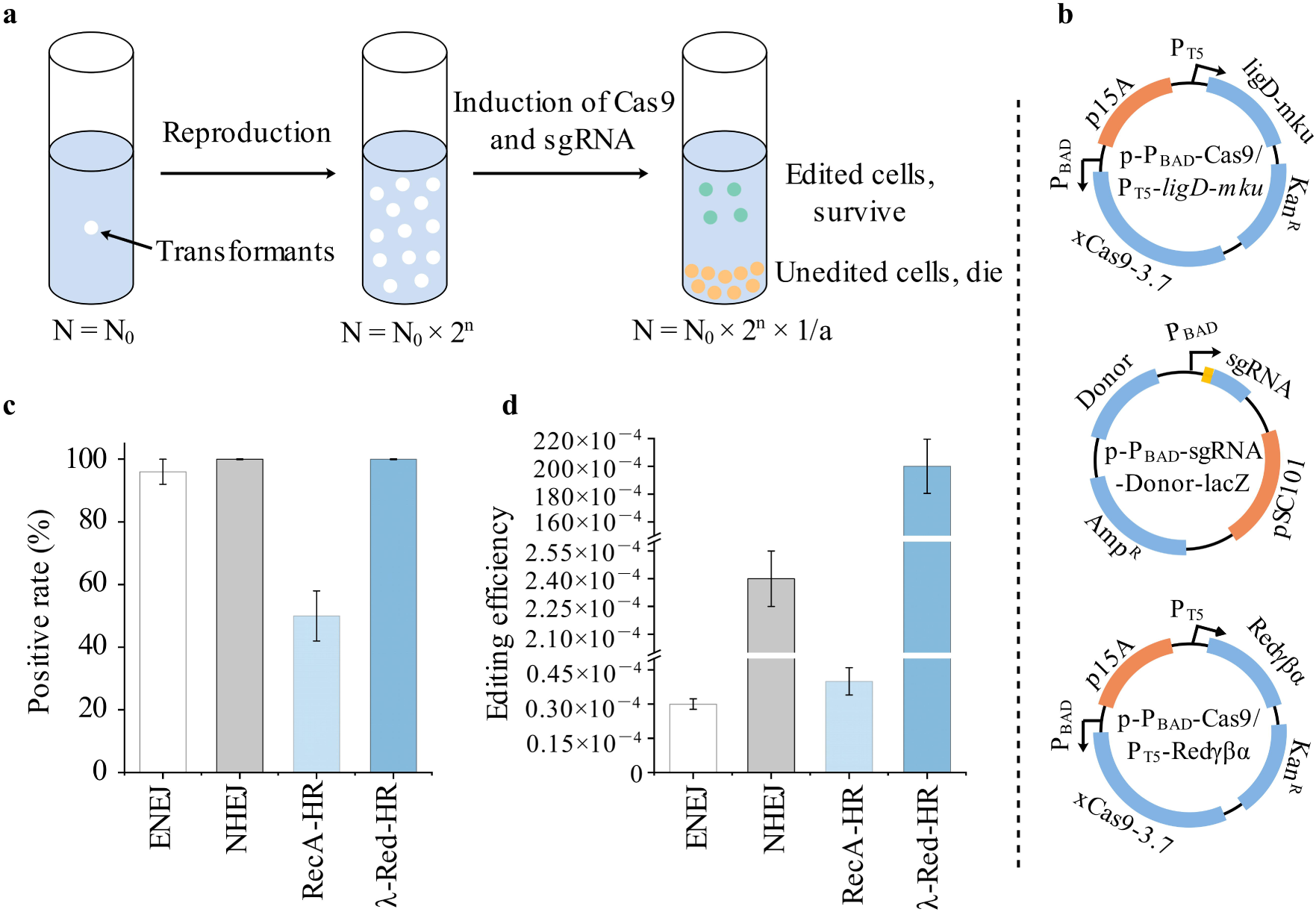
Extended methods of CNEE. (**a**) The core strategy of CNEE. Transformants are cultured for reproduction to increase the cells number and cells activity before triggering DNA cleavage and DSB repair. The white dots represent WT cells. The green dots represent edited cells. The orange dots represent cells that die from the cleavage of Cas9. “N” represents the number of living cells. “N_0_” represents the number of transformants. “n” represents the number of cell division. “1/a” represents the efficiency of DSB repair. (**b**) Plasmids constructed for extended methods. Plasmid p-P_BAD_-*cas9*/P_T5_-*ligD*-*mku* was obtained by adding the NHEJ associated genes (*ligD* and *mku*) to plasmid p-P_BAD_-*cas9*. Plasmid p-P_BAD_-*cas9*/P_T5_-Redγβα was obtained by adding λ-Red system (Redγβα) to plasmid p-P_BAD_-*cas9*. Plasmid p-P_BAD_-sgRNA-Donor-*lacZ* was obtained by adding a donor sequence to plasmid p-P_BAD_-sgRNA-*lacZ*. (**c**-**d**) Positive rates and editing efficiencies of CNEE and three extended methods. In extended methods, NHEJ pathway, RecA-mediated HR pathway, or λ-Red-mediated HR pathway served as DSB repair pathway. Data are expressed as means ± s.d. from three independent experiments.

To explore whether A-EJ involves in the CNEE, the *lacZ* was inactivated in the MG1655 (WT), MG1655 (*ΔrecA*), and MG1655 (*ΔrecBCD*) strains. Both the MG1655 (*ΔrecA*) and MG1655 (*ΔrecBCD*) strains were obtained by CNEE. In the MG1655 (*ΔrecA*) and MG1655 (*ΔligB*) strains, the positive rate were 92% and 95%, respevtively, and the editing efficiency were 3.2 × 10^−5^ and 3.15 × 10 ^−5^, respectively, close to that of the MG1655 (WT) strain (96%, 3.0 × 10 ^−5^) (Figs. 5b and 5c), indicating that DSBs repair is independent of the RecA recombinase and LigB. However, no DSB repair event was found in the MG1655 (*ΔrecBCD*) strain (Figs. 5b and 5c), demonstrating that DSBs repair relied on the RecBCD complex. Moreover, microhomology (1-3 nt) in the joining site were observed in every DSB repair event (Fig. 5d). These results agreed with the A-EJ model. We were unable to verify the role of LigA in DSB repair via gene deletion, as the *ligA* is an essential gene for *E. coli*. Taken together, the DSB repair is independent of RecA and LigB, but is dependent on RecBCD complex. Therefore, the A-EJ was likely the ENEJ involved in the CNEE.

### Strategies used in CNEE are also applicable for other CRISPR-Cas9-mediated genome editing methods

Accumulating the offsprings of the transformants harboring the editing systems and recovering the targeted cells are the keys to ensure high efficient CNEE. This strategy was applied to other CRISPR-Cas9-mediated methods depending on RecA mediated HR, λ-Red mediated HR, or NHEJ. The *Mycobacterium tuberculosis* H37Rv-derived NHEJ pathway (Mt-NHEJ) is efficient in joining non-homologous DNA ends (1). The adjacent genes *ligD* and *mku* are functional in Mt-NHEJ, and these were added to plasmid p-P_BAD_-*cas9* with the IPTG-induced P_T5_ promoter to generate plasmid p-P_BAD_-*cas9*/P_T5_-*ligD*-*mku* (Fig. 6b). The newly constructed plasmid was then transformed into MG1655 (WT) strain along with plasmid p-P_BAD_-sg*RNA*-*lacZ*. IPTG was used to induce the expression of Ligase-D and Ku protein. Other than this, all procedures were the same as CNEE. We obtained a 100% positive rate (Fig. 6c) and a 2.4 × 10^−4^ editing efficiency (Fig. 6d). The editing efficiency was eight-fold as high as that of CNEE, indicating that the Mt-NHEJ was more efficient than ENEJ in repairing CRISPR-Cas9-induced DSBs.

Most reported methods used HR to repair CRISPR-Cas9-induced DSBs. However, according to previous reports, the RecA-mediated HR pathway in *E. coli* showed low efficiency in repairing DSBs, thus, λ-Red system must be introduced to enhance HR activity in these study (6,20). Now that the ENEJ, a weak DSB repair mechanism, worked well in CNEE, it was interesting to know how RecA-mediated HR performs using the same strategies of CNEE. We added a donor sequence containing two homologous arms of 500 bp to plasmid p-P_BAD_-sg*RNA*-*lacZ* to generate plasmid p-P_BAD_-sgRNA-Donor-*lacZ* (Fig. 6b). The plasmid was transformed into MG1655 (WT) strain along with plasmid p-P_BAD_-*cas9*. All procedures were the same as CNEE. As a result, a 50% positive rate and a 4 × 10^−5^ editing efficiency were obtained (Fig. 6c). The editing efficiency was slightly higher than CNEE, and the positive rate was lower than CNEE. We found that nearly all the remaining colonies (50%) that were not positive went through DSB repair by ENEJ mechanism, indicating that RecA-mediated HR was equivalent to ENEJ in repairing CRISPR-Cas9-induced DSBs. To enhance HR activity, we added λ-Red system to plasmid p-P_BAD_-*cas9* to generate plasmid p-P_BAD_-*cas9*/*P*_T5_-*Redγβα* (Fig. 6b). The plasmid was transformed into MG1655 (WT) strain, along with plasmid p-P_BAD_-sgRNA-Donor-*lacZ*. The positive rate was 100% (Fig. 6c), indicating that all colonies grown in the plate were edited. The editing efficiency was 2 × 10^−2^, about 667-fold of CNEE, indicating that 1 out of 50 cells was edited.

## Discussion

In this study, we developed the CNEE for genomic editing. The method utilizes ENEJ to repair DSBs, thus is independent of exogenous DNA repair systems such as NHEJ. To our knowledge, this is the first successful engineering effort to utilize ENEJ mechanism in genomic editing. Accumulating the offsprings of the transformants harboring the editing systems and recovering the targeted cells are the keys to ensure the high efficiency of the CNEE. The method has been applied for gene inactivation and large-fragment knocking-out. For gene inactivation, the positive rates were above 96%, and the editing efficiencies were above 2.2 × 10 ^−5^. For large-fragment knocking-out, the positive rates were above 85%, and the editing efficiencies were above 2.6 × 10^−6^. Our method has three advantages compared to the existing methods: (i) it requires neither a DNA template nor an exogenous DSBs repair system such as NHEJ, (ii) it is simple and efficient in gene inactivation and large-fragment knocking-out, (iii) it is independent of high-efficiency competent cells.

Classic recombineering in *E. coli* relies on λ-Red recombinases to achieve genomic editing with an introduced homologous DNA as the editing template (Table 3). Either double-stranded DNA (dsDNA) or single-stranded DNA (ssDNA) is functional as the editing template. Generally, dsDNA mediated recombineering is less efficient, and it requires a selection marker such as an antibiotic resistance gene (29–31). On the contrary, ssDNA mediated recombineering does not require selection marker due to its higher efficiency; however, it is only applicable to small-scale genomic editing such as codon displacement (32,33). Recombineering is unable to knock out large fragments from the chromosome, which limits its application in large-scale genetic engineering. The introduction of CRISPR-Cas9 to *E. coli* provides an efficient way for complicated chromosome modifications such as knocking out large fragments (Table 3). Many CRISPR-Cas9-dependent methods have been reported. In these methods, the CRISPR-Cas9 system was combined with an exogenous DSB repair system, λ-Red system, or NHEJ system to achieve site-specific genomic editing. However, our testing showed that these methods are strictly dependent on the exogenous DSB repair system. In CNEE, ENEJ is employed to repair CRISPR-Cas9-induced DSBs in the absence of an editing template. The ENEJ mechanism naturally exists in *E. coli*. As a much weaker DSB repair mechanism than λ-Red system or NHEJ system, it still shows a good performance in genomic editing. By gene disruption, we demonstrated that the ENEJ is independent of RecA recombinase and LigB, but dependent on RecBCD, a protein complex that has 5’ to 3’ exonuclease, 3’ to 5’ exonuclease, and helicase activities. Theoretically, ligase is involved in the end joining of DNA ends. In *E. coli*, LigA and LigB are the only ligases. Now that the LigB is useless in DSB repair, the LigA is likely the ligase involved in the ENEJ mechanism. We were unable to verify the role of LigA in end joining because the *ligA* is an essential gene in *E. coli*. The sequencing results of over thirty samples of DSB repair indicated that microhomology of 1–3 nt was essential for end joining. The A-EJ model proposed by Chayot et al. explains the end-joining mechanism well, but the exact biochemical mechanism remains to be explored.

CRISPR-Cas9 system has been conveniently employed to introduce site-specific DSBs, and the cleavage of dsDNA kills WT cells, thereby facilitating the selection of edited cells. In eukaryotes such as *Saccharomyces cerevisiae*, the HR and NHEJ activities are high enough to efficiently repair CRISPR-Cas9-induced DSBs. Therefore, the application of CRISPR-Cas9 in eukaryotes is simple and convenient. However, in bacteria, the situation is very different. A small fraction of bacteria has relatively high HR activity such as *Streptococcus pneumoniae* or relatively high NHEJ activity such as *Mycobacterium tuberculosis*. In many bacteria such as *E. coli*, both HR and end-joining activities are weak. The application of CRISPR-Cas9 in these bacteria is, thus, difficult. However, in this study, we show that the *E. coli* native HR pathway or end-joining pathway is applicable for efficient genomic editing, which makes genomic editing in *E. coli* as simple as that in *S. cerevisiae*. This observation implies that all bacteria have the potential to be edited, rely on their native DSB repair mechanism.

Existing genomic editing methods for *E. coli* are strictly dependent on high-efficiency competent cells, and electroporation or expensive commercial kits are generally used to obtain high-efficiency competent cells. However, it is difficult to make highly competent cells for some *E. coli* strains even using special and complicated methods. Therefore, genomic editing in these strains is highly limited. However, in this study, we showed that CNEE is independent of high-efficiency competent cells, and the required minimal transformation efficiency is 10^4^ cfu/μg DNA, which is very easy to obtain using traditional, chemical methods. Moreover, the same editing efficiency is achieved as long as the transformation efficiency is over 10^4^ cfu/μg DNA. We believe that CNEE is more advantageous in other bacteria that have a lower competency than *E. coli*.

Gene inactivation and large-fragment knocking-out are frequently employed in synthetic biology and metabolic engineering. And researchers always choose to utilize an efficient and simple method. As opposed to the indirect selection provided by recombineering, the use of the CRISPR-Cas9 system makes it possible to directly select for the desired mutation. CRISPR-Cas9-induced DNA cleavage kills WT cells, and thus edited cells can be selected easily. However, the major limitation is the presence of a background of cells that escape CRISPR-Cas9-induced cell death and lack the desired mutation, and the limitation is more prominent in strains that have a weak DSB repair mechanism. Improving the effieicncy of CRISPR/Cas9 system is a key way to solve this problem. In this study, the Cas9 derivative xCas9-3.7 and the engineered sgRNA were utilized to successfully increase the efficiency of introducing DSBs (7,8). Moreover, the engineering of an inducible system helps circumvent this limitation. Off-target is another limitation for the application of CRISPR/Cas9 technology. In this study, the use of xCas9-3.7 helps to reduce the off-target event. In addition, designing an appropriate sgRNA sequence is important to avoid off-target. Therefore, a reliable sgRNA designing software or website is needed.

## Materials and methods

### Strains and culture conditions

The *E. coli* strain DH5α was used as the host strain for molecular cloning and manipulation of plasmids. The MG1655 (WT), MG1655 (*recA-*), or MG1655 (*ΔrecBCD*) were employed as host strains for genetic engineering. Strains MG1655 (*recA-*) and MG1655 (*ΔrecBCD*) were obtained by the CNEE (proposed in this study) in the basis of MG1655 (WT). Luria-Bertani (LB) medium (10 g/L tryptone, 5 g/L yeast extract, and 10 g/L NaCl) was used for cell growth in all cases unless otherwise noted. Agar was added at 20 g/L for solid medium. SOC medium (20 g/L tryptone, 5 g/L yeast extract, 0.5 g/L NaCl, 2.5 mM KCl, 10 mM MgCl2, 10 mM MgSO4, and 20 mM glucose) was used for cell recovery. Ampicillin (Amp), kanamycin (Kan) and chloramphenicol (Cm) were used at concentrations of 0.1 g/L, 0.05 g/L and 0.025 g/L, respectively. When appropriate, IPTG, X-gal, L-arabinose, and glucose were supplemented into media or cultures as follows: IPTG (1 mM), X-gal (0.1 g/L), L-arabinose (20 mM), and glucose (10 g/L).

### Plasmids construction

The primers and plasmids used in this study are listed in Tables S2 and S3, respectively. Detailed construction procedures for plasmids p-P_BAD_-sgRNA-X are shown in Fig. S11. The complete sequences of the plasmids _p-s_gRNA-template A, _p-s_gRNA-template B, _p-s_gRNA-template C, p-P_BAD_-*cas9*, p-P_BAD_-*cas9*/P_T5_-*ligD*-*mku*, and p-P_BAD_-*cas9*/P_T5_-Redγβα are presented in Notes S1-S6. The CRISPR target sequences used in this study are listed in Table S4.

### CNEE procedure

First, the Kan-resistant (Kan^R^) plasmid p-P_BAD_-*cas9* was transformed into the target strain such as MG1655 to obtain the corresponding transformant, such as MG1655/p-P_BAD_-*cas9*. A series of temperature-sensitive Amp-resistant (Amp^R^) plasmids containing specifically designed spacers were constructed to express the corresponding sgRNA and were generally named as p-P_BAD_-sgRNA-X. Then, specific plasmid p-P_BAD_-sgRNA-X was transformed into the MG1655/p-P_BAD_-*cas9* strain, and the MG1655/p-P_BAD_-*cas9*/p-P_BAD_-sgRNA-X strain was screened in a LB plate with Amp, Kan, and glucose at 30°C. A single colony was picked and inoculated into 0.5 mL SOC medium and cultured at 30°C for 2 h. Then, 4.5 mL LB medium, 5 μL Amp, and 5 μL Kan were added to the cultures. After 1 h, 100 μL of L-arabinose was added, and the cultures were cultured for another 3 h before plating. A 10-μL aliquot of the cultures were plated onto a LB plate containing Amp, Kan, and L-arabinose, and the plate was cultured overnight at 30°C. A flowchart of CNEE is shown in Fig. 2 and Fig. S9. Details of the reagents and media used in the CNEE are listed in Table S5.

### Plasmids curing and iterative editing

Positive mutants were verified by colony PCR and sequencing and were cultured in LB media individually in the presence of only Kan at 37°C for 12 h to remove the temperature-sensitive Amp^R^ plasmid p-P_BAD_-sgRNA-X (Fig. S10a). Then, the obtained edited strain containing only the plasmid p-P_BAD_-*cas9* could was used as the starting strain for the next round of genomic editing. The Kan^R^ plasmid p-P_BAD_-*cas9* is not stable in the host strain in the absence of Kan. When the last round of genomic editing was completed, the edited strain was cultured in LB media without Kan at 37°C for 16 h to remove both Amp^R^ plasmids p-P_BAD_-sgRNA-X and Kan^R^ plasmid p-P_BAD_-*cas9* (Fig. S10b). The overnight culture was diluted and plated on a LB plate. Strains sensitive to both Amp and Kan are plasmid-free. Single plasmid-free colonies were selected and verified using LB media with or without corresponding antibiotics. The flowchart of the procedure is shown in Fig. S9b.

### Measurement of transformation efficiency

Pure, supercoiled pUC19 was used to determine the transformation efficiency of competent cells. First, 1 μl of pUC19 (1 ng/μl) was added to one tube of competent cells. Next, the mixture was incubated for 30 minutes before conducting heat-shock for 90 seconds in a 42°C water bath. Then, the tube was placed on ice for two minutes before adding 1 mL of SOC medium, and the tube was shaked at 200–230 rpm (37°C) for 40 minutes. Last, 100 μl of the cultures was plated on a LB plate containing ampicillin, and the plate was incubated overnight at 37°. The transformation efficiency is N × 10^4^ cfu/μg pUC19 (“N” refers to the number of colonies).

### Calculation of genomic editing positive rate

One hundred colonies in the LB plate containing Amp, Kan, and L-arabinose were tested by colony PCR to screen for positive mutants. Twenty of the positive mutants were sequenced for further verification. The positive rate was calculated by the proportion of positive colonies to the total number of colonies.

### Calculation of genomic editing efficiency

In previous reports, the positive rate was frequently taken as the editing efficiency in a genomic editing. In this study, we proposed a new definition that better reflected editing efficiency. One control group was set along with the experimental group to calculate the editing efficiency. In the control group, L-arabinose was not added to the cultures, thus no Cas9 protein was expressed. In addition, all other operations were the same as the experimental group. The editing efficiency was calculated by the proportion of positive colonies in the experimental group to the total number of colonies in the control group.

## Acknowledgements

The work finished in Beijing Institute of Technology was supported by the National Natural Science Foundation of China (grant No. 21676026), the National Key R&D Program of China (grant No. 2017YFD0201400), the China Postdoctoral Science Foundation (grant No. 2017M620643) and the Key Program of National Natural Science Foundation of China (grant No. 21736002).

## References

1. Su T, et al. (2016) A CRISPR-Cas9 assisted non-homologous end-joining strategy for one-step engineering of bacterial genome. Sci Rep 6:37895.

2. Gu P, et al. (2015) A rapid and reliable strategy for chromosomal integration of gene(s) with multiple copies. Sci Rep 5:9684.

3. Smanski MJ, et al. (2016) Synthetic biology to access and expand nature’s chemical diversity. Nat Rev Microbiol 14:135–149.

4. Esvelt KM, Wang HH (2013) Genome-scale engineering for systems and synthetic biology. Mol Syst Biol 9:641.

5. Thomas G, Gersbach CA, Barbas CF (2013) ZFN, TALEN, and CRISPR/Cas-based methods for genome engineering. Trends Biotechnol 31:397–405.

6. Wenyan J, David B, David C, Feng Z, Marraffini LA (2013) RNA-guided editing of bacterial genomes using CRISPR-Cas systems. Nat Biotechnol 31:233–239.

7. Hu JH, et al. (2018) Evolved Cas9 variants with broad PAM compatibility and high DNA specificity. Nature 556:57–63.

8. Jinek M, et al. (2012) A programmable dual-RNA–guided DNA endonuclease in adaptive bacterial immunity. Science 337:816–821.

9. René D, et al. (2004) Evidence that stable retroviral transduction and cell survival following DNA integration depend on components of the nonhomologous end joining repair pathway. J Virol 78:8573–8581.

10. Claire W, Roland K (2006) DNA Double-strand break repair: all’s well that ends well. Annu Rev Genet 40:363–383.

11. Romain C, Benjamin M, Didier M, Miria RJ (2010) An end-joining repair mechanism in *Escherichia coli*. Proc Natl Acad Sci U.S.A. 107:2141–2146.

12. Wilson TE, Topper LM, Palmbos PL (2003) Non-homologous end-joining: bacteria join the chromosome breakdance. Trends Biochem Sci 28:62–66.

13. Marina D, et al. (2004) Mycobacterial Ku and ligase proteins constitute a two-component NHEJ repair machine. Science 306:683–685.

14. Weller GR, et al. (2002) Identification of a DNA nonhomologous end-joining complex in bacteria. Science 297:1686–1689.

15. Malyarchuk S, et al. (2007) Expression of *Mycobacterium tuberculosis* Ku and Ligase D in *Escherichia coli* results in RecA and RecB-independent DNA end-joining at regions of microhomology. DNA Repair 6:1413–1424.

16. Li Y, et al. (2015) Metabolic engineering of *Escherichia coli* using CRISPR–Cas9 meditated genome editing. Metab Eng 31:13–21.

17. Pines G, et al. (2015) Codon compression algorithms for saturation mutagenesis. ACS Synth Biol 4:604–614.

18. Zheng X, Li SY, Zhao GP, Wang J (2017) An efficient system for deletion of large DNA fragments in *Escherichia coli* via introduction of both Cas9 and the non-homologous end joining system from *Mycobacterium smegmatis*. Biochem Biophys Res 485(4):768–774.

19. Cui L, Bikard D (2016) Consequences of Cas9 cleavage in the chromosome of *Escherichia coli*. Nucleic Acids Res 44:4243–4251.

20. Zhao D, et al. (2016) Development of a fast and easy method for *Escherichia coli* genome editing with CRISPR/Cas9. Microb Cell Fact 15(1):205.

21. Cho S, et al. (2018) High-level dCas9 expression induces abnormal cell morphology in *Escherichia coli*. ACS Synth Biol 7:1085–1094.

22. Datsenko KA, Wanner BL (2000) One-step inactivation of chromosomal genes in *Escherichia coli* K-12 using PCR products. Proc Natl Acad Sci U.S.A. 97:6640–6645.

23. Baba T, et al. (2006) Construction of *Escherichia coli* K-12 in-frame, single-gene knockout mutants: the Keio collection. Mol Syst Biol 2:8.

24. Yu J, et al. (2015) Multigene editing in the *Escherichia coli* genome via the CRISPR-Cas9 system. Appl Environ Microbiol 81(7):2506–2514.

25. Zhang H, Cheng QX, Liu AM, Zhao GP, Wang J (2017) A novel and efficient method for bacteria genome editing employing both CRISPR/Cas9 and an antibiotic resistance cassette. Front Microbiol 8:812.

26. Zhao D, et al. (2017) CRISPR/Cas9-assisted gRNA-free one-step genome editing with no sequence limitations and improved targeting efficiency. Sci Rep 7(1):16624.

27. Sambrook J, Russell DW (2006) The inoue method for preparation and transformation of competent *E. Coli*: “ultra-competent” cells. CSH Protoc 2006:418–424.

28. Li J, et al. (2018) T5 exonuclease-dependent assembly offers a low-cost method for efficient cloning and site-directed mutagenesis. Nucleic Acids Res 47:3.

29. Pines G, Freed EF, Winkler JD, Gill RT (2015) Bacterial recombineering: genome engineering via phage-based homologous recombination. ACS Synth Biol 4:1176–1185.

30. Santos CN, Regitsky DD, Yoshikuni YJ (2013) Implementation of stable and complex biological systems through recombinase-assisted genome engineering. Nat Commun 4:2503.

31. Hailong W, et al. (2014) Improved seamless mutagenesis by recombineering using *ccdB* for counterselection. Nucleic Acids Res 42:e37.

32. Wang HH, et al. (2009) Programming cells by multiplex genome engineering and accelerated evolution. Nature 460:894–898.

33. Ronda C, Pedersen LE, Sommer MO, Nielsen AT (2016) CRMAGE: CRISPR optimized MAGE recombineering. Sci Rep 6:19452.

34. Yu D, et al. (2000) An efficient recombination system for chromosome engineering in *Escherichia coli*. Proc Natl Acad Sci U.S.A. 97(11):5978–5983.

35. Ellis HM, Yu D, Ditizio T, Court DL (2001) High efficiency mutagenesis, repair, and engineering of chromosomal DNA using single-stranded oligonucleotides. Proc Natl Acad Sci U.S.A. 98:6742–6746.

