## Supplementary data for "CRISPR-Cas9-assisted native end-joining editing offers a simple strategy for efficient genetic engineering in *Escherichia coli*"

### Supplementary Contents


Supplementary Figure 1

Supplementary Figure 2

Supplementary Figure 3

Supplementary Figure 4

Supplementary Figure 5

Supplementary Figure 6

Supplementary Figure 7

Supplementary Figure 8

Supplementary Figure 9

Supplementary Figure 10

Supplementary Figure 11

Supplementary Table 1

Supplementary Table 2

Supplementary Table 3

Supplementary Table 4

Supplementary Table 5

Supplementary Note 1

Supplementary Note 2

Supplementary Note 3

Supplementary Note 4

Supplementary Note 5

Supplementary Note 6

**Fig. S1**

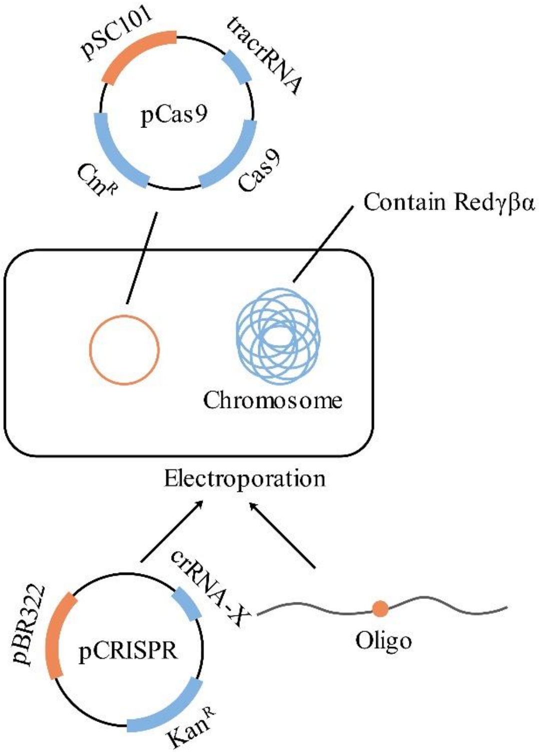


**Fig. S1.** Procedures of genomic editing of Method 1. Modified from Fig. 5 of "RNA-guided editing of bacterial genomes using CRISPR-Cas systems (Jiang *et al*., 2013)"

**Fig. S2**

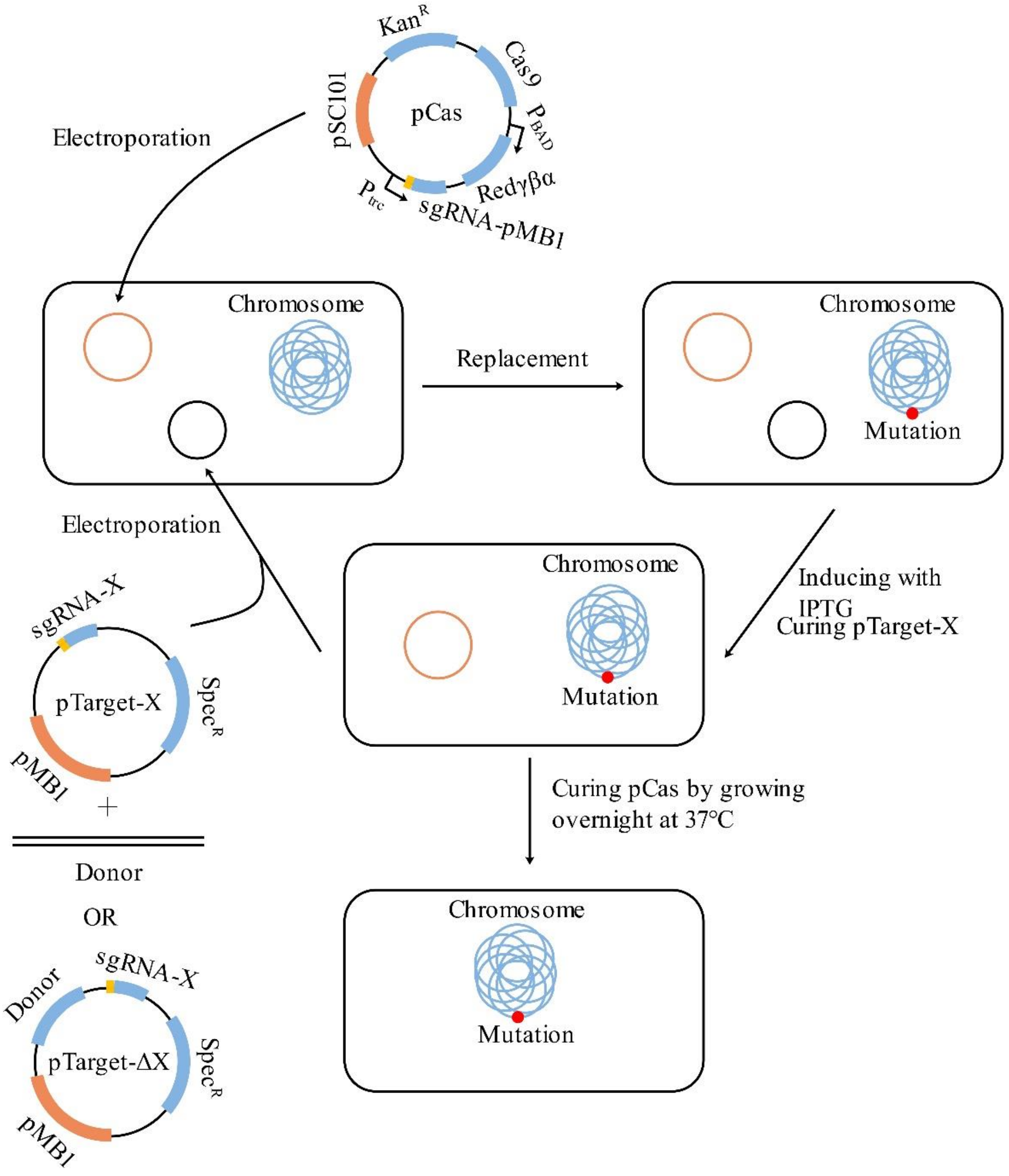


**Fig. S2.** Procedures of genomic editing of Method 2. Modified from Fig. 3 of "Multigene editing in the *Escherichia coli* genome via the CRISPR-Cas9 system (Jiang *et al*., 2015)"

**Fig. S3**

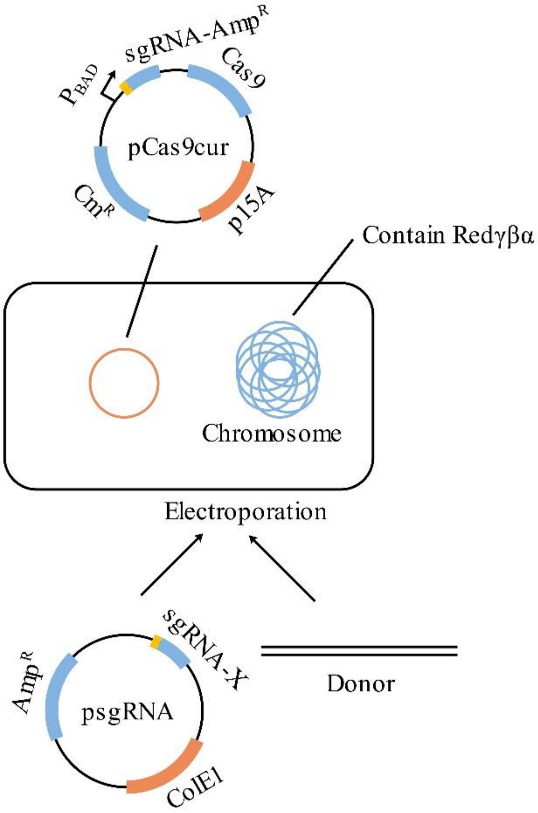


**Fig. S3.** Procedures of genomic editing of Method 3. Modified from Fig. 1 of "Metabolic engineering of *Escherichia coli* using CRISPR–Cas9 meditated genome editing (Li *et al*., 2015)"

**Fig. S4**

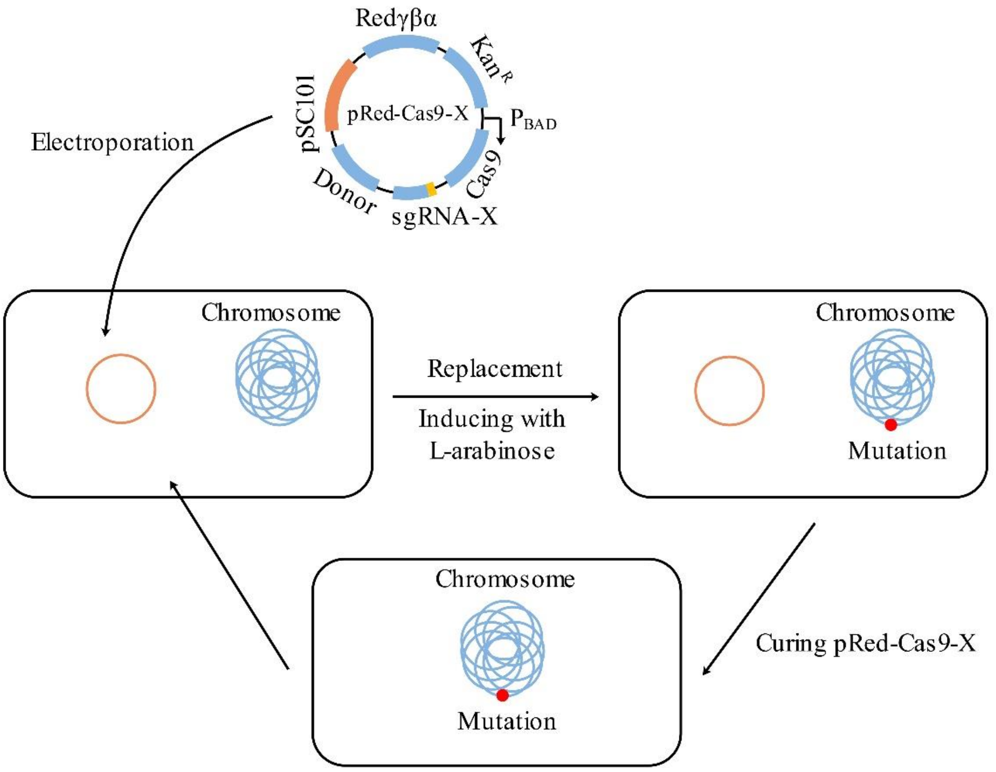


**Fig. S4.** Procedures of genomic editing of Method 4. Modified from Fig. 1 of "Development of a fast and easy method for *Escherichia coli* genome editing with CRISPR/Cas9 (Zhao *et al*., 2016)"

**Fig. S5**

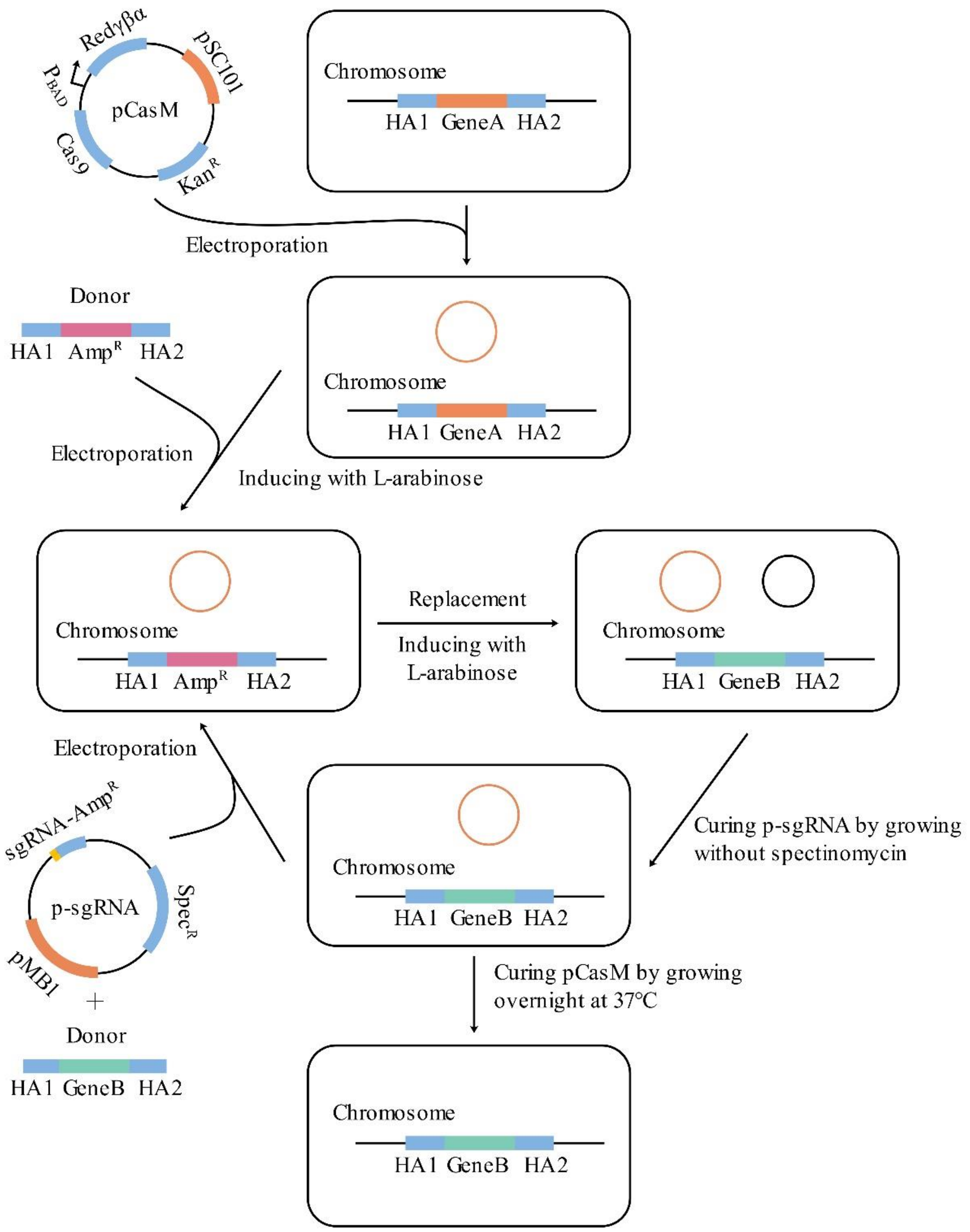


**Fig. S5.** Procedures of genomic editing of Method 5. Modified from Fig. 5 of "A novel and efficient method for bacteria genome editing employing both CRISPR/Cas9 and an antibiotic resistance cassette (Zhang *et al*., 2017)"

**Fig. S6**

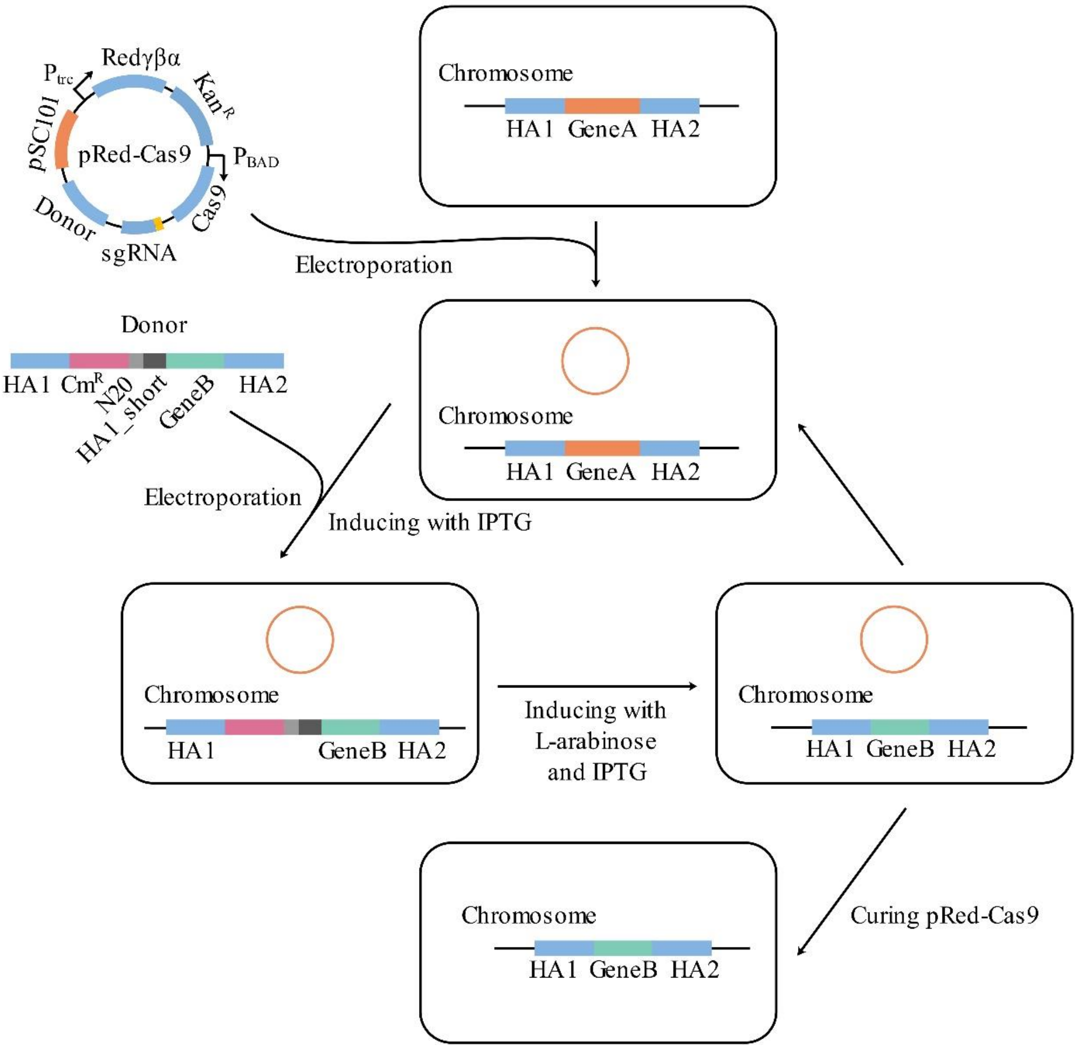


**Fig. S6.** Procedures of genomic editing of Method 6. Modified from Fig. 1 of "CRISPR/Cas9-assisted gRNA-free one-step genome editing with no sequence limitations and improved targeting efficiency (Zhao *et al.*,2017)"

**Fig. S7**

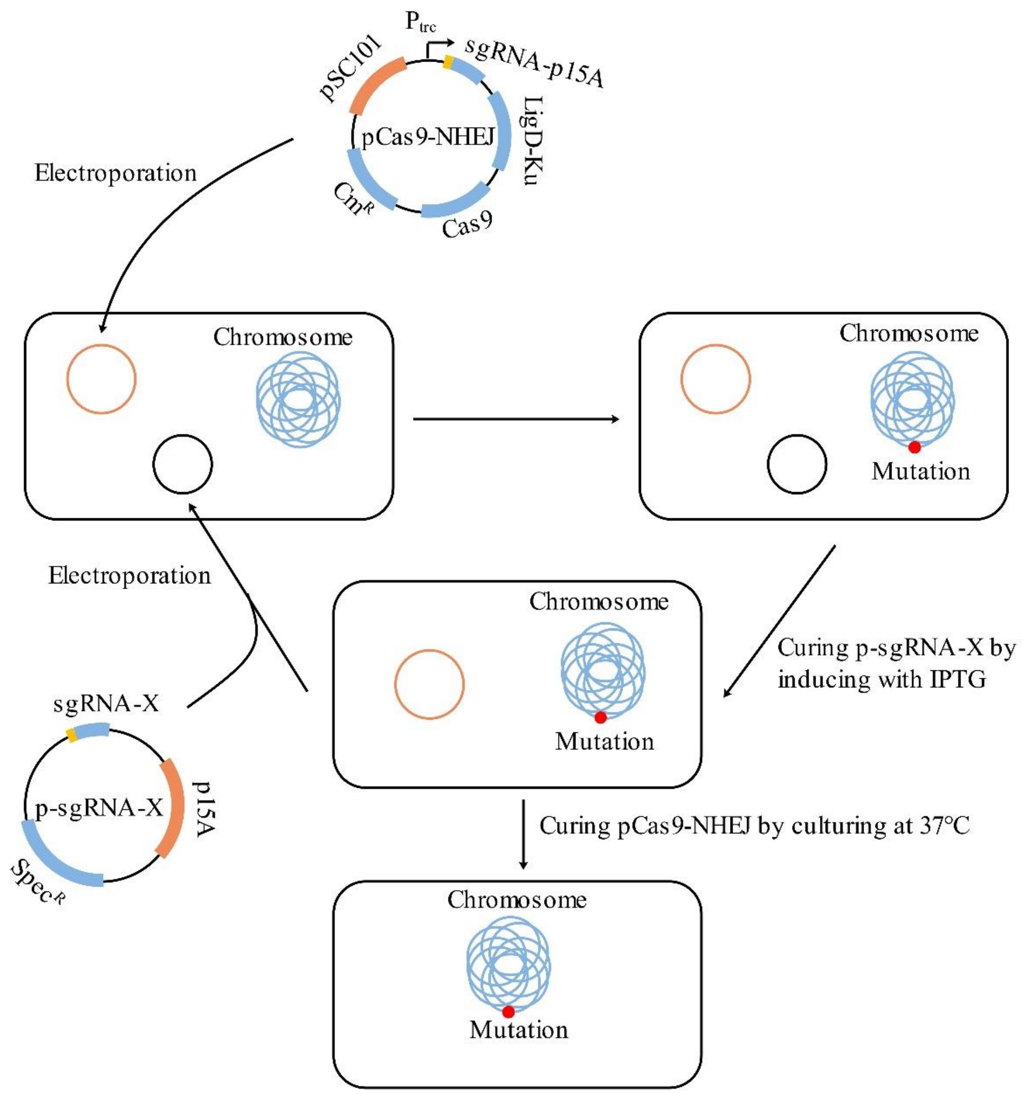


**Fig. S7.** Procedures of genomic editing of Method 7. Modified from Fig. 1 of "A CRISPR-Cas9 assisted non-homologous end-joining strategy for one-step engineering of bacterial genome (Su et al., 2016)"

**Fig. S8**

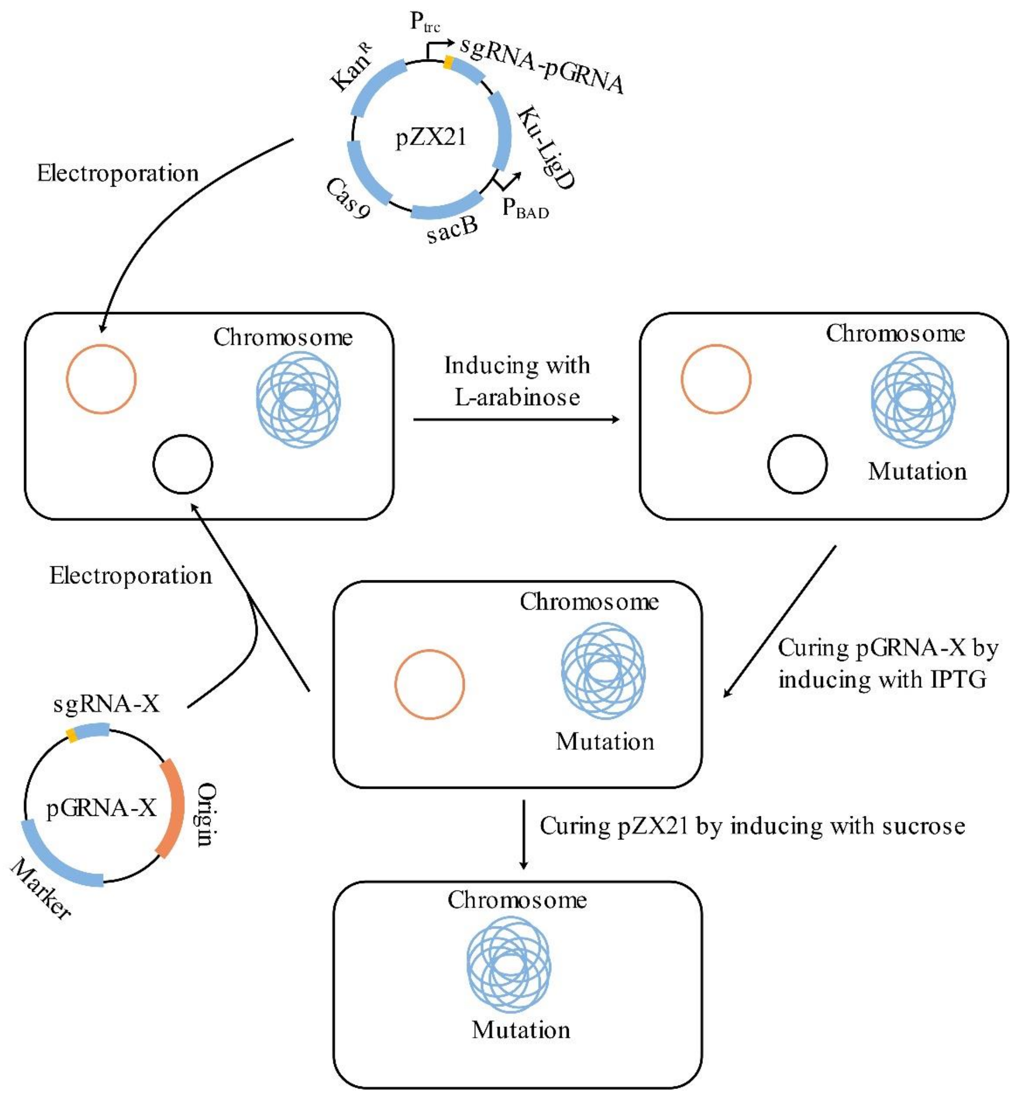


**Fig. S8.** Procedures of genomic editing of Method 8. Modified from Fig. 1 of “An efficient system for deletion of large DNA fragments in *Escherichia coli* via introduction of both Cas9 and the non-homologous end joining system from *Mycobacterium smegmatis* (Zheng., 2017)”

### Fig. S9

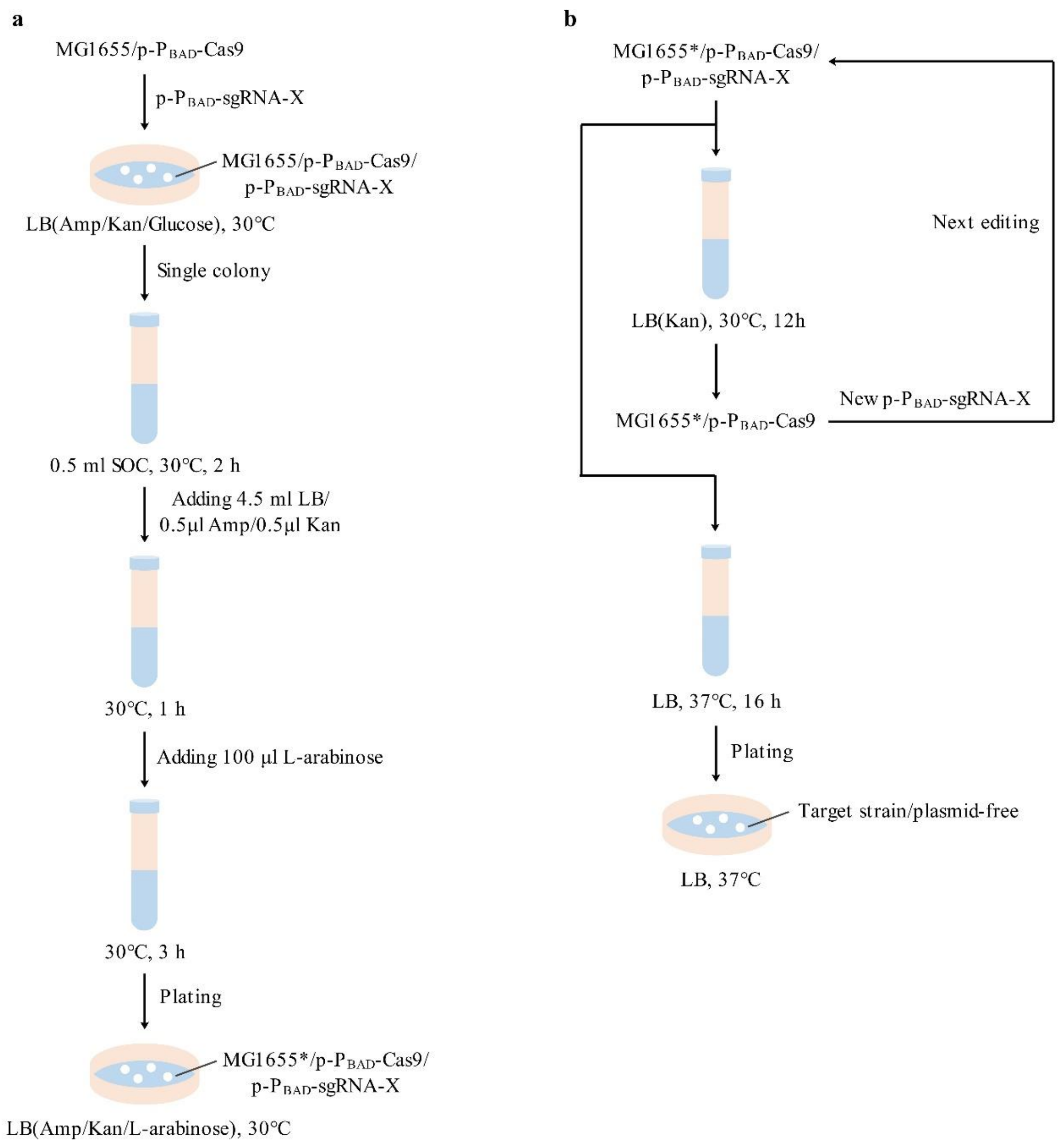


**Fig. S9.** (**a**) Detailed procedures of genomic editing. (**b**) Detailed procedures of plasmid curing and iterative editing.

**Fig. S10**

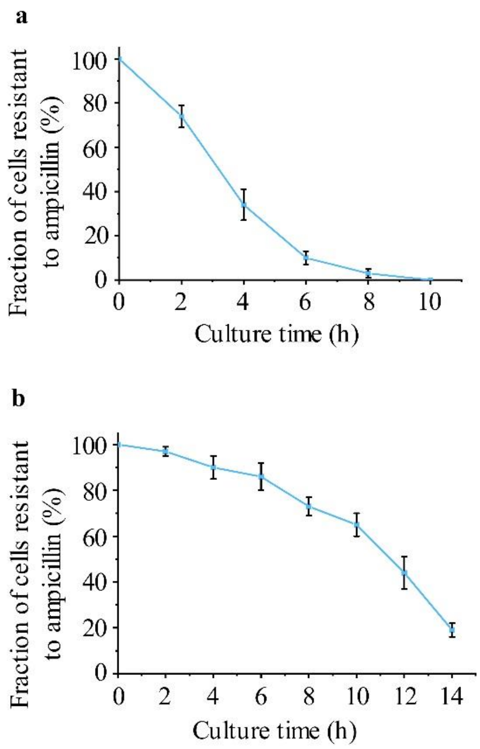


**Fig. S10.** Results of plasmid curing. (a) Plasmid curing of p-P$_{BAD}$-sgRNA-X. A single colony was inoculated in LB medium containing only kanamycin. Cells were cultivated at 37 ℃ for different period before plating on kanamycin plates. The resulting colonies were spotted on ampicillin plates to test the loss of plasmid p-P$_{BAD}$-sgRNA-X. (**b**) Plasmid curing of p-P$_{BAD}$-*cas9*. A single colony was inoculated in LB medium containing no antibiotic. Cells were cultivated at 37 ℃ for different period before plating on antibiotic-free plates. The resulting colonies were spotted on kanamycin plates to test the loss of plasmid p-P$_{BAD}$-*cas9*. Data are expressed as means ± s.d. from three independent experiments.

**Fig. S11**

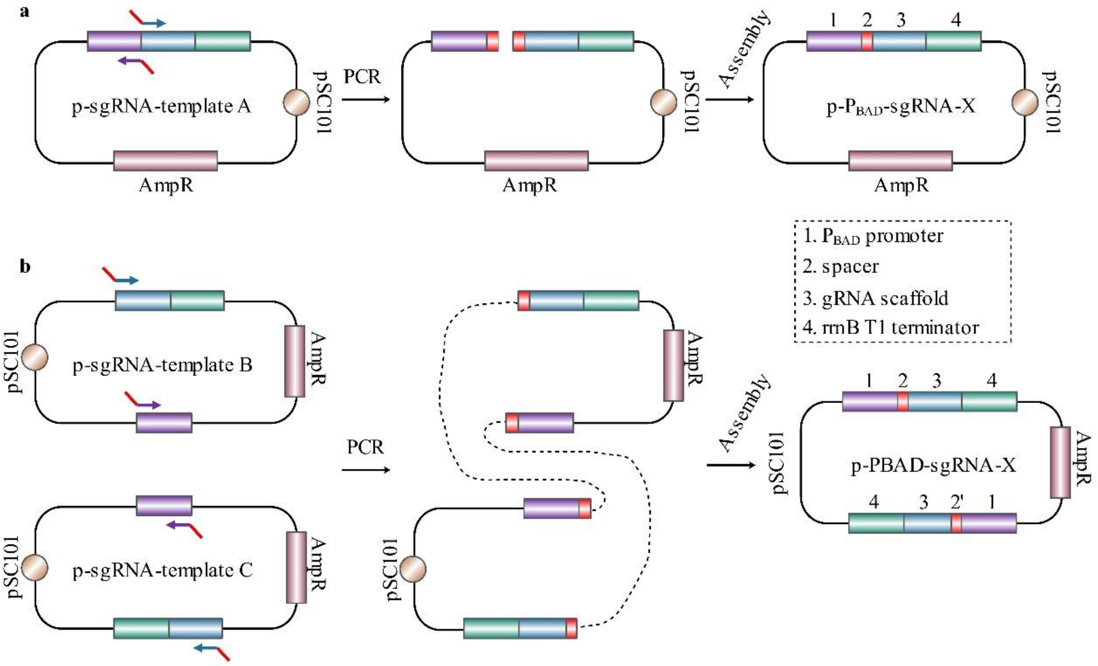


**Fig. S11.** Construction procedures of plasmid p-$P_{BAD}$-sgRNA-X. (**a**) Construction of plasmid p-$P_{BAD}$-sgRNA-X containing one sgRNA expression chimera. The plasmid p-$P_{BAD}$-template A served as a parental plasmid, and one specific spacer (20 bp) was inserted into it between the $P_{BAD}$ promoter and the gRNA scaffold via single PCR and single Gibson Assembly. The spacer introduced by PCR served as the overlap in the Gibson Assembly. (**b**) Construction of plasmid p-$P_{BAD}$-sgRNA-X containing two sgRNA expression chimeras. The plasmids p-$P_{BAD}$-template B and p-$P_{BAD}$-template C served as parental plasmids. Two fragments were obtained from the parental plasmids by PCR. Then, the two fragments were combined together via Gibson Assembly.

### Table S1. Summary of optimized conditions for CNEE

| Terms | Units | Tested conditions | Optimal condition |
|---|---|---|---|
| Inducible promoter | — | $P_{T5}$, $P_L lacO_1$, $P_{BAD}$ | $P_{BAD}$ |
| Medium 1 | — | LB, TB, SOC, SOB | SOC |
| Medium 2 | — | LB, TB, SOB | LB |
| Time 1 | hour | 0.5, 1, 1.5, 2, 2.5, 3 | 2 |
| Time 2 | hour | 0.5, 1, 1.5, 2, 2.5, 3 | 1 |
| Time 3 | hour | 1, 2, 3, 4, 5, 6 | 3 |
| L-arabinose | mM | 1, 5, 10, 15, 20, 25, 30 | 20 |
| Temperature | ℃ | 25, 28, 30, 32, 35, 37 | 30 |

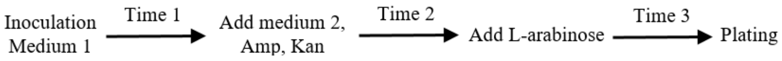

### Table S2. Primers used in this study

| Primer name | Nucleotide sequence (5'-3') |
|---|---|
| F1 | CTGACTGGCGGTTAAATTGC |
| R1 | TAACCGTCACGAGCATCATC |
| F2 | GGCCTGCCCGGTTATTATTA |
| R2 | CAGCTATGACCATGATTACGG |
| F3 | ATGCGTAAAATCATTCATGTGGA |
| R3 | TCATAATCCCAGCACCAGTTG |
| F4 | GCCGCTTATTTCTATTCCGG |
| R4 | CTGCTTACACTCTTCGGCAA |
| F5 | CCATTCCATGTCAAACCCCT |
| R5 | CAGCTCTGAACTTCTTCTGGT |
| F6 | TTCCGCCGCATGTGGATTA |
| R6 | GCACAACTTGACGTGGTGA |
| F7 | GAACGTTTTCTCGCAAACTCA |
| R7 | CCAGTGTTCAGCCATAAAGG |
| F8 | CCGCCTTCCTCTCTTCTATT |
| R8 | CCTCATACGGTTCGACGTTT |
| F9 | GTTCACCGCATCGTTAAGCA |
| R9 | TTACCACTTCATCCCGATGC |

### Table S3. Plasmids used in this study

| Plasmid name | Description | Reference |
|---|---|---|
| pCas9cur | Acquiring p15A origin, SpCas9 | Lab stock |
| pKD46 | Acquiring λ-Redγβα, $P_{BAD}$ promoter, *araC*, pSC101 origin and *rep101* | Lab stock |
| pCPB-37-441 | Acquiring $P_{T5}$ promoter | Lab stock |
| pUC19 | Acquiring $Amp^{R}$; measuring transformation efficiency | Lab stock |
| pYK-J23100-*ligD-mku* | Acquiring $Kan^{R}$, *lacI* and *ligD-mku* | Lab stock |
| pDTT-30 | Acquiring gRNA scaffold and rrnB T1 terminator | Lab stock |
| p-$P_{BAD}$-*cas9* | One of the two plasmids in the CAEE system | This study |
| p-$P_{BAD}$-*cas9*/$P_{T5}$-ligD-mku | p-$P_{BAD}$-*cas9* containing $P_{T5}$-*ligD-mku* | This study |
| p-$P_{BAD}$-*cas9*/$P_{T5}$-Redγβα | p-$P_{BAD}$-*cas9* containing $P_{T5}$-Redγβα | This study |
| p-sgRNA-template A | Construct plasmid p-$P_{BAD}$-sgRNA-X containing one spacer | This study |
| p-sgRNA-template B | Construct plasmid p-$P_{BAD}$-sgRNA-X containing two spacers | This study |
| p-sgRNA-template C | Construct plasmid p-$P_{BAD}$-sgRNA-X containing two spacers | This study |
| p-$P_{BAD}$-sgRNA-*lacZ* | Inactivate the *lacZ* gene | This study |
| p-$P_{BAD}$-sgRNA-Donor-*lacZ* | Inactivate the *lacZ* gene | This study |
| p-$P_{BAD}$-sgRNA-*dinB* | Inactivate the *dinB* gene | This study |
| p-$P_{BAD}$-sgRNA-*recA* | Inactivate the *recA* gene | This study |
| p-$P_{BAD}$-sgRNA-*recBCD* | Inactivate the *recBCD* genes | This study |
| p-$P_{BAD}$-sgRNA-83kb | Delete the 83 kb chromosome fragment | This study |
| p-$P_{BAD}$-sgRNA-81kb | Delete the 81 kb chromosome fragment | This study |

### **Table S4.** CRISPR target sequences used in this study

| Spacer | Nucleotide sequence (5’-3’) |
|---|---|
| *lacZ* | CAGTATCCCCGTTTACAGGG |
| *dinB* | ACGCCGGGGATTTTTGCCAG |
| *recA* | GAAAGAGGGCGAAAACGTGG |
| recBCD-L | GCGAGAGCTGGTTTATACCG |
| recBCD-R | ACGCGCGAAGGTAACCCCGG |
| 81 kb-L | TCCGCAGCAAAATCAGGGGG |
| 81 kb-R | GCCACATCCACTTTTTCCGG |
| 83 kb-L | TGGTGACTGAAGATATGGCG |
| 83 kb-R | ATCTATGGTGATGATGCCGG |

### Table S5. Reagents and media used in the CAEE

| Reagent | Formulation concentration | Application concentration |
|---|---|---|
| Ampicillin | 100 g/L | 0.1 g/L |
| Kanamycin | 50 g/L | 0.05 g/L |
| Glucose | 500 g/L | 10 g/L |
| L-arabinose | 1 M | 20 mM |
| Media | Composition | |
| LB | 10 g/L tryptone, 5 g/L yeast extract, 10 g/L NaCl; Solid medium (20 g/L Agar) | |
| SOC | 20 g/L tryptone, 5 g/L yeast extract, 0.5 g/L NaCl, 2.5 mM KCl, 10 mM MgCl2, 10 mM MgSO4, 20 mM glucose | |

### Note S1. Complete sequence of the plasmid p-sgRNA-template A in Genbank format

```
LOCUS       p-sgRNA-template A      3893 bp    DNA     circular SYN 24-NOV-2018
DEFINITION  p-sgRNA-template A
ACCESSION   p-sgRNA-template A
KEYWORDS
SOURCE
  ORGANISM

FEATURES             Location/Qualifiers
     CDS             complement(1..951)
                     /label="rep101"
                     /note="replication protein for the pSC101 origin"
     rep_origin      999..1221
                     /label="pSC101 ori"
                     /note=" replication origin"
     CDS             complement(1843..2703)
                     /label="AmpR"
                     /note="ampicillin resistance gene"
     promoter        complement(2704..2808)
                     /label="AmpR promoter"
     misc_feature    3230..3247
                     /label="I-SceI"
                     /note="recognition sequence of homing endonuclease I-SceI"
     promoter        3393..3558
                     /label="araBAD promoter"
     misc_RNA        3559..3634
                     /label="gRNA scaffold"
                     /note="guide RNA scaffold for the CRISPR/Cas9 system"
     terminator      3695..3766
                     /label="rrnB T1 terminator"
ORIGIN
        1 tcagatcctt ccgtatttag ccagtatgtt ctctagtgtg gttcgttgtt tttgcgtgag
       61 ccatgagaac gaaccattga gatcatactt actttgcatg tcactcaaaa attttgcctc
      121 aaaactggtg agctgaattt ttgcagttaa agcatcgtgt agtgtttttc ttagtccgtt
      181 acgtaggtag gaatctgatg taatggttgt tggtattttg tcaccattca tttttatctg
      241 gttgttctca agttcggtta cgagatccat ttgtctatct agttcaactt ggaaaatcaa
      301 cgtatcagtc gggcggcctc gcttatcaac caccaatttc atattgctgt aagtgtttaa
      361 atctttactt attggtttca aaacccattg gttaagcctt ttaaactcat ggtagttatt
      421 ttcaagcatt aacatgaact taaattcatc aaggctaatc tctatatttg ccttgtgagt
      481 tttcttttgt gttagttctt ttaataacca ctcataaatc ctcatagagt atttgttttc
      541 aaaagactta acatgttcca gattatattt tatgaatttt tttaactgga aaagataagg
```

601 caatatctct tcactaaaaa ctaattctaa tttttcgctt gagaacttgg catagtttgt
661 ccactggaaa atctcaaagc ctttaaccaa aggattcctg atttccacag ttctcgtcat
721 cagctctctg gttgctttag ctaatacacc ataagcattt tccctactga tgttcatcat
781 ctgagcgtat tggttataag tgaacgatac cgtccgttct ttccttgtag ggttttcaat
841 cgtggggttg agtagtgcca cacagcataa aattagcttg gtttcatgct ccgttaagtc
901 atagcgacta atcgctagtt catttgcttt gaaaacaact aattcagaca tacatctcaa
961 ttggtctagg tgattttaat cactatacca attgagatgg gctagtcaat gataattact
1021 agtccttttc ctttgagttg tgggtatctg taaattctgc tagacctttg ctggaaaact
1081 tgtaaattct gctagaccct ctgtaaattc cgctagacct ttgtgtgttt tttttgttta
1141 tattcaagtg gttataattt atagaataaa gaaagaataa aaaaagataa aaagaataga
1201 tcccagccct gtgtataact cactacttta gtcagttccg cagtattaca aaaggatgtc
1261 gcaaacgctg tttgctcctc tacaaaacag accttaaaac cctaaaggct taagtagcac
1321 cctcgcaagc tcggttgcgg ccgcaatcgg gcaaatcgct gaatattcct tttgtctccg
1381 accatcaggc acctgagtcg ctgtcttttt cgtgacattc agttcgctgc gctcacggct
1441 ctggcagtga atgggggtaa atggcactac aggcgccttt tatggattca tgcaaggaaa
1501 ctacccataa tacaagaaaa gcccgtcacg ggcttctcag ggcgttttat ggcgggtctg
1561 ctatgtggtg ctatctgact tttgctgtt cagcagttcc tgccctctga ttttccagtc
1621 tgaccacttc ggattatccc gtgacaggtc attcagactg gctaatgcac ccagtaaggc
1681 agcggtatca tcaacggggt ctgacgctca gtggaacgaa aactcacgtt aagggatttt
1741 ggtcatgaga ttatcaaaaa ggatcttcac ctagatcctt ttaaattaaa aatgaagttt
1801 taaatcaatc taaagtatat atgagtaaac ttggtctgac agttaccaat gcttaatcag
1861 tgaggcacct atctcagcga tctgtctatt tcgttcatcc atagttgcct gactccccgt
1921 cgtgtagata actacgatac gggagggctt accatctggc cccagtgctg caatgatacc
1981 gcgagaccca cgctcaccgg ctccagattt atcagcaata aaccagccag ccggaagggc
2041 cgagcgcaga agtggtcctg caactttatc cgcctccatc cagtctatta attgttgccg
2101 ggaagctaga gtaagtagtt cgccagttaa tagtttgcgc aacgttgttg ccattgctac
2161 aggcatcgtg gtgtcacgct cgtcgtttgg tatggcttca ttcagctccg gttcccaacg
2221 atcaaggcga gttacatgat cccccatgtt gtgcaaaaaa gcggttagct ccttcggtcc
2281 tccgatcgtt gtcagaagta agttggccgc agtgttatca ctcatggtta tggcagcact
2341 gcataattct cttactgtca tgccatccgt aagatgcttt tctgtgactg gtgagtactc
2401 aaccaagtca ttctgagaat agtgtatgcg gcgaccgagt tgctcttgcc cggcgtcaat
2461 acgggataat accgcgccac atagcagaac tttaaaagtg ctcatcattg gaaaacgttc
2521 ttcggggcga aaactctcaa ggatcttacc gctgttgaga tccagttcga tgtaacccac
2581 tcgtgcaccc aactgatctt cagcatcttt tactttcacc agcgtttctg ggtgagcaaa
2641 aacaggaagg caaaatgccg caaaaaaggg aataagggcg acacggaaat gttgaatact
2701 catactcttc ctttttcaat attattgaag catttatcag ggttattgtc tcatgagcgg
2761 atacatattt gaatgtattt agaaaaataa acaaataggg gttccgcggc acagatgcgt
2821 aaggagaaaa taccgcatca ggcgccattc gccattcagg ctgcgcaact gttgggaagg
2881 gcgatcggtg cgggcctctt cgctattacg ccagctggcg aaagggggat gtgctgcaag
2941 gcgattaagt tgggtaacgc cagggttttc ccagtcacga cgttgtaaaa cgacggccag
3001 tgccaagctt gcatgcctgc aggtcgactc tagaggatcc ccgggtaccg agctcgaatt
3061 cgtaatcatg tcatagctgt ttcctgtgtg aaattgttat ccgctcacaa ttccacacaa
3121 catacgagcc ggaagcataa agtgtaaagc ctggggtgcc taatgagtga gctaactcac

3181 attaattgcg ttgcgctcac tgcccgcttt ccagtcggga aacctgtcat agggataaca
3241 gggtaatact tttcatactc ccgccattca gagaagaaac caattgtcca tattgcatca
3301 gacattgccg tcactgcgtc ttttactggc tcttctcgct aaccaaaccg gtaaccccgc
3361 ttattaaaag cattctgtaa caaagcggga ccaaagccat gacaaaaacg cgtaacaaaa
3421 gtgtctataa tcacggcaga aaagtccaca ttgattattt gcacggcgtc acactttgct
3481 atgccatagc atttttatcc ataagattag cggatcctac ctgacgcttt ttatcgcaac
3541 tctctactgt ttctccatgt tttagagcta gaaatagcaa gttaaaataa ggctagtccg
3601 ttatcaactt gaaaaagtgg caccgagtcg gtgcttagca tccaaactcg agtaaggatc
3661 attaaggatc ccatggtacg cgtgctagag gcatcaaata aaacgaaagg ctcagtcgaa
3721 agactgggcc tttcgtttta tctgttgttt gtcggtgaac gctctcctga gtaggacaaa
3781 tccgccccat gggtatggac agttttccct ttgatatgta acggtgaaca gttgttctac
3841 ttttgtttgt tagtcttgat gcttcactga tagatacaag agccataaga acc
//

### Note S2. Complete sequence of the plasmid p-sgRNA-template B in Genbank format

```
LOCUS       p-sgRNA-template B        2998 bp        DNA        circular SYN 24-NOV-2018
DEFINITION  p-sgRNA-template B
ACCESSION   p-sgRNA-template B
KEYWORDS
SOURCE
  ORGANISM

FEATURES             Location/Qualifiers
     promoter        96..200
                     /label="AmpR promoter"
     CDS             201..1061
                     /label="AmpR"
                     /note="ampicillin resistance gene"
     promoter        1339..1504
                     /label="araBAD promoter"
     rep_origin      1539..2127
                     /label="colE1 ori"
                     /note="replication origin"
     misc_RNA        2727..2802
                     /label="gRNA scaffold"
                     /note="guide RNA scaffold for the CRISPR/Cas9 system"
     terminator      2863..2934
                     /label="rrnB T1 terminator"
ORIGIN
        1 gacgaaaggg cctcgtgata cgcctatttt tataggttaa tgtcatgata ataatggttt
       61 cttagacgtc aggtggcact tttcggggaa atgtgcgcgg aacccctatt tgtttatttt
      121 tctaaataca ttcaaatatg tatccgctca tgagacaata accctgataa atgcttcaat
      181 aatattgaaa aaggaagagt atgagtattc aacatttccg tgtcgccctt attccctttt
      241 ttgcggcatt ttgccttcct gtttttgctc acccagaaac gctggtgaaa gtaaaagatg
      301 ctgaagatca gttgggtgca cgagtgggtt acatcgaact ggatctcaac agcggtaaga
      361 tccttgagag ttttcgcccc gaagaacgtt ttccaatgat gagcactttt aaagttctgc
      421 tatgtggcgc ggtattatcc cgtattgacg ccgggcaaga gcaactcggt cgccgcatac
      481 actattctca gaatgacttg gttgagtact caccagtcac agaaaagcat cttacggatg
      541 gcatgacagt aagagaatta tgcagtgctg ccataaccat gagtgataac actgcggcca
      601 acttacttct gacaacgatc ggaggaccga aggagctaac cgcttttttg cacaacatgg
      661 gggatcatgt aactcgcctt gatcgttggg aaccggagct gaatgaagcc ataccaaacg
      721 acgagcgtga caccacgatg cctgtagcaa tggcaacaac gttgcgcaaa ctattaactg
      781 gcgaactact tactctagct tcccggcaac aattaataga ctggatggag gcggataaag
      841 ttgcaggacc acttctgcgc tcggcccttc cggctggctg gtttattgct gataaatctg
      901 gagccggtga gcgtgggtct cgcggtatca ttgcagcact ggggccagat ggtaagccct
```

961 cccgtatcgt agttatctac acgacgggga gtcaggcaac tatggatgaa cgaaatagac
1021 agatcgctga gataggtgcc tcactgatta agcattggta actgtcagac caagtttact
1081 catatatact ttagattgat ttaaaacttc atttttaatt taaaaggatc taggtgaaga
1141 tcctttttga taatctcatg accaaaatcc cttaacgtga gttttcgttc cactttttc
1201 atactcccgc cattcagaga agaaaccaat tgtccatatt gcatcagaca ttgccgtcac
1261 tgcgtctttt actggctctt ctcgctaacc aaaccggtaa ccccgcttat taaaagcatt
1321 ctgtaacaaa gcgggaccaa agccatgaca aaaacgcgta acaaaagtgt ctataatcac
1381 ggcagaaaag tccacattga ttatttgcac ggcgtcacac tttgctatgc catagcattt
1441 ttatccataa gattagcgga tcctacctga cgctttttat cgcaactctc tactgtttct
1501 ccatcgtcag accccgtaga aaagatcaaa ggatcttctt gagatccttt ttttctgcgc
1561 gtaatctgct gcttgcaaac aaaaaaacca ccgctaccag cggtggtttg tttgccggat
1621 caagagctac caactctttt tccgaaggta actggcttca gcagagcgca gataccaaat
1681 actgtccttc tagtgtagcc gtagttaggc caccacttca agaactctgt agcaccgcct
1741 acatacctcg ctctgctaat cctgttacca gtggctgctg ccagtggcga taagtcgtgt
1801 cttaccgggt tggactcaag acgatagtta ccggataagg cgcagcggtc gggctgaacg
1861 gggggttcgt gcacacagcc cagcttggag cgaacgacct acaccgaact gagataccta
1921 cagcgtgagc tatgagaaag cgccacgctt cccgaaggga gaaaggcgga caggtatccg
1981 gtaagcggca gggtcggaac aggagagcgc acgagggagc ttccaggggg aaacgcctgg
2041 tatctttata gtcctgtcgg gtttcgccac ctctgacttg agcgtcgatt tttgtgatgc
2101 tcgtcagggg ggcggagcct atggaaaaac gccagcaacg cggccttttt acggttcctg
2161 gccttttgct ggccttttgc tcacatgttc tttcctgcgt tatcccctga ttctgtggat
2221 aaccgtatta ccgcctttga gtgagctgat accgctgcac agatgcgtaa ggagaaaata
2281 ccgcatcagg cgccattcgc cattcaggct gcgcaactgt tgggaagggc gatcggtgcg
2341 ggcctcttcg ctattacgcc agctggcgaa agggggatgt gctgcaaggc gattaagttg
2401 ggtaacgcca gggttttccc agtcacgacg ttgtaaaacg acggccagtg ccaagcttgc
2461 atgcctgcag gtcgactcta gaggatcccc gggtaccgag ctcgaattcg taatcatgtc
2521 atagctgttt cctgtgtgaa attgttatcc gctcacaatt ccacacaaca tacgagccgg
2581 aagcataaag tgtaaagcct ggggtgccta atgagtgagc taactcacat taattgcgtt
2641 gcgctcactg cccgctttcc agtcgggaaa cctgtcataa caccgtgcgt gttgactatt
2701 ttacctctgg cggtgataat ggttgcgttt tagagctaga aatagcaagt taaaataagg
2761 ctagtccgtt atcaacttga aaaagtggca ccgagtcggt gcttagcatc caaactcgag
2821 taaggatcat taaggatccc atggtacgcg tgctagaggc atcaaataaa acgaaaggct
2881 cagtcgaaag actgggcctt tcgttttatc tgttgtttgt cggtgaacgc tctcctgagt
2941 aggacaaatc cgccctgcat gtgtcagagg ttttcaccgt catcaccgaa acgcgcga
//

### Note S3. Complete sequence of the plasmid p-sgRNA-template C in Genbank format

```
LOCUS       p-sgRNA-template C      3942 bp     DNA     circular SYN 24-NOV-2018
DEFINITION  p-sgRNA-template C
ACCESSION   p-sgRNA-template C
KEYWORDS
SOURCE
  ORGANISM

FEATURES             Location/Qualifiers
     CDS             complement(1..951)
                     /label="rep101"
                     /note="=replication protein for the pSC101 origin"
     rep_origin      999..1221
                     /label="pSC101 ori"
                     /note=" replication origin"
     misc_feature    1843..1860
                     /label="I-SceI"
                     /note="recognition sequence of homing endonuclease I-SceI"
     promoter        2006..2171
                     /label="araBAD promoter"
     CDS             complement(2172..3032)
                     /label="AmpR"
                     /note="ampicillin resistance gene"
     promoter        complement(3033..3137)
                     /label="AmpR promoter"
     misc_RNA        3608..3683
                     /label="gRNA scaffold"
                     /note="guide RNA scaffold for the CRISPR/Cas9 system"
     terminator      3744..3815
                     /label="rrnB T1 terminator"
ORIGIN
        1 tcagatcctt ccgtatttag ccagtatgtt ctctagtgtg gttcgttgtt tttgcgtgag
       61 ccatgagaac gaaccattga gatcatactt actttgcatg tcactcaaaa attttgcctc
      121 aaaactggtg agctgaattt ttgcagttaa agcatcgtgt agtgtttttc ttagtccgtt
      181 acgtaggtag gaatctgatg taatggttgt tggtattttg tcaccattca tttttatctg
      241 gttgttctca agttcggtta cgagatccat ttgtctatct agttcaactt ggaaaatcaa
      301 cgtatcagtc gggcggcctc gcttatcaac caccaatttc atattgctgt aagtgtttaa
      361 atctttactt attggtttca aaacccattg gttaagcctt ttaaactcat ggtagttatt
      421 ttcaagcatt aacatgaact taaattcatc aaggctaatc tctatatttg ccttgtgagt
      481 tttcttttgt gttagttctt ttaataacca ctcataaatc ctcatagagt atttgttttc
      541 aaaagactta acatgttcca gattatattt tatgaatttt tttaactgga aaagataagg
```

601 caatatctct tcactaaaaa ctaattctaa tttttcgctt gagaacttgg catagtttgt
661 ccactggaaa atctcaaagc ctttaaccaa aggattcctg atttccacag ttctcgtcat
721 cagctctctg gttgctttag ctaatacacc ataagcattt tccctactga tgttcatcat
781 ctgagcgtat tggttataag tgaacgatac cgtccgttct ttccttgtag ggttttcaat
841 cgtggggttg agtagtgcca cacagcataa aattagcttg gtttcatgct ccgttaagtc
901 atagcgacta atcgctagtt catttgcttt gaaaacaact aattcagaca tacatctcaa
961 ttggtctagg tgattttaat cactatacca attgagatgg gctagtcaat gataattact
1021 agtccttttc ctttgagttg tgggtatctg taaattctgc tagacctttg ctggaaaact
1081 tgtaaattct gctagaccct ctgtaaattc cgctagacct ttgtgtgttt tttttgttta
1141 tattcaagtg gttataattt atagaataaa gaaagaataa aaaaagataa aaagaataga
1201 tcccagccct gtgtataact cactacttta gtcagttccg cagtattaca aaaggatgtc
1261 gcaaacgctg tttgctcctc tacaaaacag accttaaaac cctaaaggct taagtagcac
1321 cctcgcaagc tcggttgcgg ccgcaatcgg gcaaatcgct gaatattcct tttgtctccg
1381 accatcaggc acctgagtcg ctgtcttttt cgtgacattc agttcgctgc gctcacggct
1441 ctggcagtga atgggggtaa atggcactac aggcgccttt tatggattca tgcaaggaaa
1501 ctacccataa tacaagaaaa gcccgtcacg ggcttctcag ggcgttttat ggcgggtctg
1561 ctatgtggtg ctatctgact ttttgctgtt cagcagttcc tgccctctga ttttccagtc
1621 tgaccacttc ggattatccc gtgacaggtc attcagactg gctaatgcac ccagtaaggc
1681 agcggtatca tcaacggggt ctgacgctca gtggaacgaa aactcacgtt aagggatttt
1741 ggtcatgaga ttatcaaaaa ggatcttcac ctagatcctt ttaaattaaa aatgaagttt
1801 taaatcaatc taaagtatat atgagtaaac ttggtctgac agtagggata acagggtaat
1861 acttttcata ctcccgccat tcagagaaga aaccaattgt ccatattgca tcagacattg
1921 ccgtcactgc gtcttttact ggctcttctc gctaaccaaa ccggtaaccc cgcttattaa
1981 aagcattctg taacaaagcg ggaccaaagc catgacaaaa acgcgtaaca aaagtgtcta
2041 taatcacggc agaaaagtcc acattgatta tttgcacggc gtcacacttt gctatgccat
2101 agcattttta tccataagat tagcggatcc tacctgacgc tttttatcgc aactctctac
2161 tgtttctcca tttaccaatg cttaatcagt gaggcaccta tctcagcgat ctgtctattt
2221 cgttcatcca tagttgcctg actccccgtc gtgtagataa ctacgatacg ggagggctta
2281 ccatctggcc ccagtgctgc aatgataccg cgagacccac gctcaccggc tccagattta
2341 tcagcaataa accagccagc cggaagggcc gagcgcagaa gtggtcctgc aactttatcc
2401 gcctccatcc agtctattaa ttgttgccgg gaagctagag taagtagttc gccagttaat
2461 agtttgcgca acgttgttgc cattgctaca ggcatcgtgg tgtcacgctc gtcgtttggt
2521 atggcttcat tcagctccgg ttcccaacga tcaaggcgag ttacatgatc ccccatgttg
2581 tgcaaaaaag cggttagctc cttcggtcct ccgatcgttg tcagaagtaa gttggccgca
2641 gtgttatcac tcatggttat ggcagcactg cataattctc ttactgtcat gccatccgta
2701 agatgctttt ctgtgactgg tgagtactca accaagtcat tctgagaata gtgtatgcgg
2761 cgaccgagtt gctcttgccc ggcgtcaata cgggataata ccgcgccaca tagcagaact
2821 ttaaaagtgc tcatcattgg aaaacgttct tcggggcgaa aactctcaag gatcttaccg
2881 ctgttgagat ccagttcgat gtaacccact cgtgcaccca actgatcttc agcatctttt
2941 actttcacca gcgtttctgg gtgagcaaaa acaggaaggc aaaatgccgc aaaaaaggga
3001 ataagggcga cacggaaatg ttgaatactc atactcttcc tttttcaata ttattgaagc
3061 atttatcagg gttattgtct catgagcgga tacatatttg aatgtattta gaaaaataaa
3121 caaatagggg ttccgcggca cagatgcgta aggagaaaat accgcatcag gcgccattcg

3181 ccattcaggc tgcgcaactg ttgggaaggg cgatcggtgc gggcctcttc gctattacgc
3241 cagctggcga aagggggatg tgctgcaagg cgattaagtt gggtaacgcc agggttttcc
3301 cagtcacgac gttgtaaaac gacggccagt gccaagcttg catgcctgca ggtcgactct
3361 agaggatccc cgggtaccga gctcgaattc gtaatcatgt catagctgtt tcctgtgtga
3421 aattgttatc cgctcacaat tccacacaac atacgagccg gaagcataaa gtgtaaagcc
3481 tggggtgcct aatgagtgag ctaactcaca ttaattgcgt tgcgctcact gcccgctttc
3541 cagtcgggaa acctgtcata acaccgtgcg tgttgactat tttacctctg gcggtgataa
3601 tggttgcgtt ttagagctag aaatagcaag ttaaaataag gctagtccgt tatcaacttg
3661 aaaaagtggc accgagtcgg tgcttagcat ccaaactcga gtaaggatca ttaaggatcc
3721 catggtacgc gtgctagagg catcaaataa aacgaaaggc tcagtcgaaa gactgggcct
3781 ttcgttttat ctgttgtttg tcggtgaacg ctctcctgag taggacaaat ccgccccatg
3841 ggtatggaca gttttccctt tgatatgtaa cggtgaacag ttgttctact tttgtttgtt
3901 agtcttgatg cttcactgat agatacaaga gccataagaa cc

//

### Note S4. Complete sequence of the plasmid p-$P_{BAD}$-*cas9* in Genbank format

LOCUS p-$P_{BAD}$-*cas9* 9354 bp DNA circular SYN 24-NOV-2018
DEFINITION p-$P_{BAD}$-*cas9*
ACCESSION p-$P_{BAD}$-*cas9*
KEYWORDS
SOURCE
ORGANISM

FEATURES Location/Qualifiers
promoter 1745..1849
/label="AmpR promoter"
CDS 618..1412
/label=KanR
/note="aminoglycoside phosphotransferase from Tn5"
CDS complement(2822..6928)
/label="xCas9-3.7"
/note="generates RNA-guided double strand breaks in DNA"
promoter complement(6972..7137)
/label="araBAD promoter"
/note="promoter of the L-arabinose operon of E. coli"
CDS 7283..8161
/label="araC"
/note="L-arabinose regulatory protein"
terminator 8205..8251
/label="rrn B T1 terminator"
rep_origin complement(8583..9128)
/label="p15A ori"
/note="replication origin"
ORIGIN
1 aacttttata acaaataatc aaggagaaat tcaaagaaat ttatcagccg tgtcgccctt
61 aattgtgagc ggataacaat tacgagcttc atgcacagtg aaatcatgaa aaatttattt
121 gctttgtgag cggataacaa ttataatatg tggaattgtg agcgctcaca attccacaac
181 ggtttccctc tagaaataat tttgtttaac ttttcgagac cttaggaggt aaacatcgca
241 tcctcacgat aatatccggg taggcgcaat cactttcgtc tactccgtta caaagcgagg
301 ctgggtattt cccggccttt ctgttatccg aaatccactg aaagcacagc ggctggctga
361 ggagataaat aataaacgag ggctgtatg cacaaagcat cttctgttga gttaagaacg
421 agtatcgaga tggcacatag ccttgctcaa attggaatca ggtttgtgcc aataccagta
481 gaaacagacg aagaatccat gggtatggac agcgcggaac ccctatttgt ttatttttct
541 aaatacattc aaatatgtat ccgctcatga gacaataacc ctgataaatg cttcaataat
601 attgaaaaag gaagagtatg attgaacaag atggattgca cgcaggttct ccggccgctt
661 gggtggagag gctattcggc tatgactggg cacaacagac aatcggctgc tctgatgccg

721 ccgtgttccg gctgtcagcg caggggcgcc cggttctttt tgtcaagacc gacctgtccg
781 gtgccctgaa tgaactgcag gacgaggcag cgcggctatc gtggctggcc acgacgggcg
841 ttccttgcgc agctgtgctc gacgttgtca ctgaagcggg aagggactgg ctgctattgg
901 gcgaagtgcc ggggcaggat ctcctgtcat ctcaccttgc tcctgccgag aaagtatcca
961 tcatggctga tgcaatgcgg cggctgcata cgcttgatcc ggctacctgc ccattcgacc
1021 accaagcgaa acatcgcatc gagcgagcac gtactcggat ggaagccggt cttgtcgatc
1081 aggatgatct ggacgaagag catcaggggc tcgcgccagc cgaactgttc gccaggctca
1141 aggcgcgcat gcccgacggc gaggatctcg tcgtgaccca tggcgatgcc tgcttgccga
1201 atatcatggt ggaaaatggc cgcttttctg gattcatcga ctgtggccgg ctgggtgtgg
1261 cggaccgcta tcaggacata gcgttggcta cccgtgatat tgctgaagag cttggcggcg
1321 aatgggctga ccgcttcctc gtgctttacg gtatcgccgc tcccgattcg cagcgcatcg
1381 ccttctatcg ccttcttgac gagttcttct gaaaggaggt tataaaaaat gaaaccagta
1441 acgttatacg atgtcgcaga gtatgccggt gtctcttatc agaccgtttc ccgcgtggtg
1501 aaccaggcca gccacgtttc tgcgaaaacg cgggaaaaag tggaagcggc gatggcggag
1561 ctgaattaca ttcccaaccg cgtggcacaa caactggcgg gcaaacagtc gttgctgatt
1621 ggcgttgcca cctccagtct ggccctgcac gcgccgtcgc aaattgtcgc ggcgattaaa
1681 tctcgcgccg atcaactggg tgccagcgtg gtggtgtcga tggtagaacg aagcggcgtc
1741 gaagcctgta aagcggcggt gcacaatctt ctcgcgcaac gcgtcagtgg gctgatcatt
1801 aactatccgc tggatgacca ggatgccatt gctgtggaag ctgcctgcac taatgttccg
1861 gcgttatttc ttgatgtctc tgaccagaca cccatcaaca gtattatttt ctcccatgaa
1921 gacggtacgc gactgggcgt ggagcatctg gtcgcattgg gtcaccagca aatcgcgctg
1981 ttagcgggcc cattaagttc tgtctcggcg cgtctgcgtc tggctggctg gcataaatat
2041 ctcactcgca atcaaattca gccgatagcg gaacgggaag gcgactggag tgccatgtcc
2101 ggttttcaac aaaccatgca aatgctgaat gagggcatcg ttcccactgc gatgctggtt
2161 gccaacgatc agatggcgct gggcgcaatg cgcgccatta ccgagtccgg gctgcgcgtt
2221 ggtgcggata tctcggtagt gggatacgac gataccgaag acagctcatg ttatatcccg
2281 ccgttaacca ccatcaaaca ggattttcgc ctgctggggc aaaccagcgt ggaccgcttg
2341 ctgcaactct ctcagggcca ggcggtgaag ggcaatcagc tgttgcccgt ctcactggtg
2401 aaaagaaaaa ccaccctggc gcccaatacg caaaccgcct ctccccgcgc gttggccgat
2461 tcattaatgc agctggcacg acaggtttcc cgactggaaa gcgggcagtg ataactgtca
2521 gaccaagttt acgagctcgc ttggactcct gttgatagat ccagtaatga cctcagaact
2581 ccatctggat ttgttcagaa cgctagggat aacagggtaa ttattgcgct gcagttttgg
2641 gaccattcaa aacagcatag ctctaaaacc tcgtagacta tttttgtcta aaaaatttcg
2701 taatcgcact atttgtctca gctagacttc agtcttgaaa agcccctgta ttactgcatt
2761 tattaagagt attataccat atttttagtt attaagaaat aatcttcatc taaaatatac
2821 ttcagtcacc tcctagctga ctcaaatcaa tgcgtgtttc ataaagacca gtgatggatt
2881 gatggataag agtggcatct aaaacttctt ttgtagacgt atatcgttta cgatcaattg
2941 ttgtatcaaa atatttaaaa gcagcgggag ctccaagatt cgtcaacgta aataaatgaa
3001 taatattttc tgcttgttca cgtattggtt tgtctctatg tttgttatat gcactaagaa
3061 ctttatctaa attggcatct gctaaaataa cacgcttaga aaattcactg atttgctcaa
3121 taatctcatc taaataatgc ttatgctgct ccacaaacaa ttgtttttgt tcgttatctt
3181 ctggactacc cttcaacttt tcataatgac tagctaaata taaaaaattc acatatttgc
3241 ttggcagagc cagctcattt ccttttgta acactccggc actagccagc atccgtttac

3301 gaccgttttc taactcaaaa agactatatt taggtagttt aatgattaag tcttttttaa
3361 cttccttata tcctttagct tctaaaaagt caatcggatt tttttcaaag gaacttcttt
3421 ccataattgt gatccctagt aactctttaa cggattttaa cttcttcgat ttcccttttt
3481 ccaccttagc aaccactagg actgaataag ctaccgttgg actatcaaaa ccaccatatt
3541 tttttggatc ccagtctttt ttacgagcaa taagcttgtc cgaatttctt tttggtaaaa
3601 ttgactcctt ggagaatccg cctgtctgta cttctgtttt cttgacaata ttgacttggg
3661 gcatggacaa tactttgcgc actgtggcaa aatctcgccc tttatcccag acaatttctc
3721 cagtttcccc attagtttcg attagagggc gtttgcgaat ctctccattt gcaagtgtaa
3781 tttctgtttt gaagaagttc atgatattag agtaaaagaa atattttgcg gttgctttgc
3841 ctatttcttg ctcagactta gcaatcattt tacgaacatc ataaacttta taatcaccat
3901 agacaaactc cgattcaagt tttggatatt tcttaatcaa agcagttcca acgacggcat
3961 ttagatacgc atcatgggca tgatggtaat tgttaatctc acgtacttta tagaattgga
4021 aatcttttcg gaagtcagaa actaatttag attttaaggt aatcacttta acctctcgaa
4081 taagtttatc attttcatcg tatttagtat tcatgcgact atccaaaatt tgtgccacat
4141 gcttagtgat ttggcgagtt tcaaccaatt ggcgtttgat aaaaccagct ttatcaagtt
4201 cactcaaacc tccacgttca gctttcgtta aattatcaaa cttacgttga gtgattaact
4261 tggcgtttag aagttgtctc caatagtttt tcatcttttt gactacttct tcacttggaa
4321 cgttatccga tttaccacga tttttatcag aacgcgttaa gaccttattg tctattgaat
4381 cgtctttaag gaaactttgt ggaacaatgt gatcgacatc ataatcactt aaacgattaa
4441 tatctaattc ttggtccaca tacatgtctc ttccattttg gagataatag agatagagct
4501 tttcattttg caattgagta ttttcaacag gatgctcttt aagaatctga cttcctaatt
4561 ctttgatacc ttcttcgatt cgtttcatac gctctcgcga atttttctgg cccttttgag
4621 ttgtctgatt ttcacgtgcc atttcaataa cgatattttc tggcttatgc cgccccatta
4681 ctttgaccaa ttcatcaaca acttttacag tctgtaaaat accttttta atagcagggc
4741 taccagctaa atttgcaata tgttcatgta aactatcgcc ttgtccagac acttgtgctt
4801 tttgaatgtc ttctttaaat gtcaaactat catcatggat cagctgaata aaattgcgat
4861 tggcaaaacc atctgatttc aaaaaatcta atattgtttt gccagattgc ttatccctaa
4921 taccattaat caattttcga gacaaacgtc cccaaccagt ataacggcga cgtttaagct
4981 gtttcatcac cttatcatca aagaggtgag catatgtttt aagtctttcc tcaatcatct
5041 ccctatcttc aaataaggtc aatgttaaaa caatatcctc taagatatct tcattttctt
5101 cattatccaa aaaatcttta tctttaataa tttttagcaa atcatggtag gtacctaatg
5161 aagcattaaa tctatcttca actcctgaaa tttcaacact atcaaaacat tctatttttt
5221 tgaaataatc ttcttttaat tgcttaacgg ttacttttcg atttgttttg aagagtaaat
5281 caacaatggc tttcttctga tcacctgaaa gaaatgctgg ttttcgcatt ccttcagtaa
5341 catatttgac ctttgtcaat tcgttataaa ccgtaaaata ctcataaagc aaactatgtt
5401 ttggtagtac tttttcattt ggaagatttt tatcaaagtt tgtcatgcgt tcaataaatg
5461 attgagctga agcaccttta tcgacaactt tttcaaaatt ccatggggta attgtttctt
5521 cagacttccg agtcatccat gcaaaacgac tattgccacg cgccaatgga ccaacataat
5581 aaggaattcg aaaagtcaag atttttttcaa tcttctcacg attgtctttt aaaaatggat
5641 aaaagtcttc ttgtcttctc aaaatagcat gcagctcacc caagtgaatt tgatggggaa
5701 taatgccgtt gtcaaaggtc cgttgcttgc gcagcaaatc ttcacgattt agtttcacca
5761 ataattcctc agtaccatcc attttttcta aaattggttt gataaattta taaaattctt
5821 cttggctagc tcccccatca atataacctg catatccgtt ttttgattga tcaaaaaaga

5881 tttctttata cttttctgga agttgttgtc gaactaaagc ttttaaaaga gtcaagtctt
5941 gatgatgttc atcgtacagt ttaatcattg aagctgatag gggagcctta gttatttcag
6001 tatttactct taggatatct gaaagtaaaa tagcatctga taaattctta gctgccaaaa
6061 acaaatcagc atattgatct ccaatttgcg ccaataaatt atctaaatca tcatcgtaag
6121 tatcttttga aagctgtaat ttggtatctt ctgccaaatc aaaatttgat ttaaaattag
6181 gggtcaaacc caatgacaaa gcaatgagat tcccaaataa gccatttttc ttctcaccgg
6241 ggagctgagc aatgagattt tctaatcgtc ttgatttact caatcgtgca gaaagaatcg
6301 ctttagcatc tactccactt gcgttaatag ggttttcttc aaataattga ttgtaggttt
6361 gtaccaactg gataaatagt ttgtccacat cactattatc aggatttaaa tctccctcaa
6421 tcaaaaaatg accacgaaac ttaatcatat gcgctaaggc caaatagatt aagcgcaaat
6481 ccgctttatc agtagaatct accaattttt ttcgcagatg atagatagtt ggatatttct
6541 catgataagc aacttcatct actatatttc caaaaatagg atgacgttca tgcttcttgt
6601 cttcttccac caaaaaagac tcttcaagtc gatgaaagaa actatcatct actttcgcca
6661 tctcatttga aaaaatctcc tgtagataac aaatacgatt cttccgacgt gtataccttc
6721 tacgagctgt ccgtttgaga cgagtcgctt ccgctgtctc tccactgtca aataaaagag
6781 cccctataag atttttttg atactgtggc ggtctgtatt tcccagaacc ttgaactttt
6841 tagacggaac cttatattca tcagtgatca ccgcccatcc gacgctattt gtgccgatat
6901 ctaagcctat tgagtatttc ttatccattt tttataacct ccttagagct cgaattccca
6961 aaaaaacggg tatggagaaa cagtagagag ttgcgataaa aagcgtcagg taggatccgc
7021 taatcttatg gataaaaatg ctatggcata gcaaagtgtg acgccgtgca aataatcaat
7081 gtggactttt ctgccgtgat tatagacact tttgttacgc gtttttgtca tggctttggt
7141 cccgctttgt tacagaatgc ttttaataag cggggttacc ggtttggtta gcgagaagag
7201 ccagtaaaag acgcagtgac ggcaatgtct gatgcaatat ggacaattgg tttcttctct
7261 gaatggcggg agtatgaaaa gtatggctga agcgcaaaat gatcccctgc tgccgggata
7321 ctcgtttaat gcccatctgg tggcgggttt aacgccgatt gaggccaacg gttatctcga
7381 ttttttatc gaccgaccgc tgggaatgaa aggttatatt ctcaatctca ccattcgcgg
7441 tcagggggtg gtgaaaaatc agggacgaga atttgtttgc cgaccgggtg atattttgct
7501 gttcccgcca ggagagattc atcactacgg tcgtcatccg gaggctcgcg aatggtatca
7561 ccagtgggtt tactttcgtc cgcgcgccta ctggcatgaa tggcttaact ggccgtcaat
7621 atttgccaat acggggttct ttcgcccgga tgaagcgcac cagccgcatt tcagcgacct
7681 gtttgggcaa atcattaacg ccgggcaagg ggaagggcgc tattcggagc tgctggcgat
7741 aaatctgctt gagcaattgt tactgcggcg catggaagcg attaacgagt cgctccatcc
7801 accgatggat aatcgggtac gcgaggcttg tcagtacatc agcgatcacc tggcagacag
7861 caattttgat atcgccagcg tcgcacagca tgtttgcttg tcgccgtcgc gtctgtcaca
7921 tcttttccgc cagcagttag ggattagcgt cttaagctgg cgcgaggacc aacgtatcag
7981 ccaggcgaag ctgcttttga gcaccacccg gatgcctatc gccaccgtcg gtcgcaatgt
8041 tggttttgac gatcaactct atttctcgcg ggtatttaaa aaatgcaccg gggccagccc
8101 gagcgagttc cgtgccggtt gtgaagaaaa agtgaatgat gtagccgtca agttgtcata
8161 ataaatcgat gcaggtggca cttttcgggg aaatgtggag gcatcaaata aaacgaaagg
8221 ctcagtcgaa agactgggcc tttcgtttta tctgttgttt gtcggtgaac gctctcctga
8281 gtaggacaaa tccgccgccc tagacctagg gcgttcggct gcggcgagcg gtatcagctc
8341 actcaaaggc ggtaatacgg ttatccacag aatcagggga taacgcagga aagagcatgt
8401 gagcaaaagg ccagcaaaag gccaggaacc gtggatatat tccgcttcct cgctcactga

8461 ctcgctacgc tcggtcgttc gactgcggcg agcggaaatg gcttacgaac ggggcggaga
8521 tttcctggaa gatgccagga agatacttaa cagggaagtg agagggccgc ggcaaagccg
8581 tttttccata ggctccgccc ccctgacaag catcacgaaa tctgacgctc aaatcagtgg
8641 tggcgaaacc cgacaggact ataaagatac caggcgtttc cccctggcgg ctccctcgtg
8701 cgctctcctg ttcctgcctt tcggtttacc ggtgtcattc cgctgttatg gccgcgtttg
8761 tctcattcca cgcctgacac tcagttccgg gtaggcagtt cgctccaagc tggactgtat
8821 gcacgaaccc cccgttcagt ccgaccgctg cgccttatcc ggtaactatc gtcttgagtc
8881 caacccggaa agacatgcaa aagcaccact ggcagcagcc actggtaatt gatttagagg
8941 agttagtctt gaagtcatgc gccggttaag gctaaactga aaggacaagt tttggtgact
9001 gcgctcctcc aagccagtta cctcggttca aagagttggt agctcagaga accttcgaaa
9061 aaccgccctg caaggcggtt ttttcgtttt cagagcaaga gattacgcgc agaccaaaac
9121 gatctcaaga agatcatctt attaataagg atctcaagaa gatcctttga tcttttctac
9181 ggggtctgac gctcagtgga acgaaaactc acgttaaggg attttggtca tgactagtgc
9241 ttggattctc accaataaaa aacgcccggc ggcaaccgat ttcaagttga taacggacta
9301 gccttatttt aacttgctat gctgttttga atggttccaa caagattatt ttat

//

### Note S5. Complete sequence of the plasmid p-$P_{BAD}$-*cas9*/$P_{T5}$-*ligD-mku* in Genbank format

LOCUS p-$P_{BAD}$-*cas9*/$P_{T5}$-*ligD-mku* 12476 bp DNA circular SYN 24-NOV-2018
DEFINITION p-$P_{BAD}$-*cas9*/$P_{T5}$-*ligD-mku*
ACCESSION p-$P_{BAD}$-*cas9*/$P_{T5}$-*ligD-mku*
KEYWORDS
SOURCE
ORGANISM

FEATURES Location/Qualifiers
promoter 1..50
/label="T5 promoter"
CDS 134..2413
/label="LigD"
CDS 2434..3255
/label="Ku"
terminator 3256..3500
/label="lambda tL3 terminator"
promoter 3532..3636
/label="AmpR promoter"
CDS 3637..4431
/label=KanR
/note="aminoglycoside phosphotransferase from Tn5"
CDS 4448..5530
/label="lacI"
/note="lac repressor"
terminator 5563..5622
/label="lamda t0 terminator"
CDS complement(5841..9947)
/label="xCas9-3.7"
/note="generates RNA-guided double strand breaks in DNA"
promoter complement(9991..10156)
/label="araBAD promoter"
/note="promoter of the L-arabinose operon of E. coli"
CDS 10302..11180
/label="araC"
/note="L-arabinose regulatory protein"
terminator 11224..11270
/label="rrn B T1 terminator"
rep_origin complement(11602..12147)
/label="p15A ori"
/note="replication origin"

ORIGIN

```
        1 tcatgaaaaa tttatttgct ttgtgagcgg ataacaatta taatatgtgg aattgtgagc
       61 gctcacaatt ccacaacggt ttccctctag aaataatttt gtttaacttt tcgagacctt
      121 aggaggtaaa catatgggtt cggcgtcgga gcaacgggtg acgctgacca acgccgacaa
      181 ggtgctctat cccgccaccg ggaccacaaa gtccgatatc ttcgactact acgccggtgt
      241 tgccgaagtc atgctcggcc acatcgcggg acggccggcg acgcgcaagc gctggcctaa
      301 cggcgtcgac caacccgcgt tcttcgaaaa gcagttggcg ttgtcggcgc cgccttggct
      361 gtcacgtgca acggtggcgc accggtccgg gacgacgacc tatccgatca tcgatagcgc
      421 aaccgggctg gcctggatcg cccaacaggc ggcgctggag gtgcacgtgc cgcagtggcg
      481 gtttgtcgcc gagcccggat caggtgagtt aaatccgggc ccggcaacgc gtttggtgtt
      541 cgacctggac ccgggcgaag gcgtgatgat ggcccagctg gccgaggtgg cgcgcgcggt
      601 tcgtgatctt ctcgccgata tcgggttggt caccttcccg gtcaccagcg gcagcaaggg
      661 attgcatctg tacacaccgc tggatgagcc ggtgagcagc aggggagcca cggtgttggc
      721 caagcgcgtc gcgcagcgat tggagcaggc gatgcccgcg ttggtcacct cgaccatgac
      781 caaaagcctg cgggccggga aggtgtttgt ggactggagc cagaacagcg gctcgaagac
      841 caccatcgcg ccgtactcac tacgtggccg gacgcatccg accgtcgcgg cgccacgcac
      901 ctgggcggag ctcgacgacc ccgcactgcg tcagctctcc tacgacgagg tgctgacccg
      961 gattgcccgc gacggcgatc tgctcgagcg gctggatgcc gacgctccgg tagcggaccg
     1021 gttgacccga taccgccgca tgcgcgacgc atcgaaaact cccgagccga ttcccacggc
     1081 gaaacccgtt accggagacg gcaatacgtt cgtcatccag gagcatcacg cgcgtcggcc
     1141 gcactacgat ttccggctgg aatgcgacgg cgtgctggtc tcgtgggcgg taccgaaaaa
     1201 cctgcccgac aacacatcgg ttaaccatct agcgatacac accgaggacc acccgctgga
     1261 atacgccacg ttcgagggcg cgattcccag cggggagtac ggcgccggca aggtgatcat
     1321 ctgggactcc ggcacttacg acaccgagaa gttccacgat gacccgcaca cgggggaggt
     1381 catcgtgaat ctgcacggcg gccggatctc tgggcgttat gcgctgattc ggaccaacgg
     1441 cgatcggtgg ctggcgcacc gcctaaagaa tcagaaagac cagaaggtgt tcgagttcga
     1501 caatctggcc ccaatgcttg ccacgcacgg cacggtggcc ggtctaaagg ccagccagtg
     1561 ggcgttcgaa ggcaagtggg acggctaccg gttgctggtt gaggctgacc acggcgccgt
     1621 gcggctgcgg tcccgcagcg ggcgcgatgt caccgccgag tatccgcaat tgcgggcatt
     1681 ggcggaggat ctcgccgatc accacgtggt gctggacggc gaggccgtcg tacttgactc
     1741 ctctggtgtg cccagcttca gccagatgca gaatcggggc cgcgacaccc gtgtcgagtt
     1801 ctgggcgttc gacctgctct acctcgacgg ccgcgcgctg ctaggcaccc gctaccaaga
     1861 ccggcgtaag ctgctcgaaa ccctagctaa cgcaaccagt ctcaccgttc ccgagctgct
     1921 gcccggtgac ggcgcccaag cgtttgcgtg ctcgcgcaag cacggctggg agggcgtgat
     1981 cgccaagagg cgtgactcgc gctatcagcc gggccggcgc tgcgcgtcgt gggtcaagga
     2041 caagcactgg aacacccagg aagtcgtcat tggtggctgg cgcgccgggg aaggcgggcg
     2101 cagcagtggc gtcgggtcgc tgctcatggg catccccggt ccaggtgggc tgcagttcgc
     2161 cgggcgggtc ggtaccggcc tcagcgaacg cgaactggcc aacctcaagg agatgctggc
     2221 gccgctgcat accgacgagt ccccctcga cgtaccactg cccgcgcgtg acgccaaggg
     2281 catcacatat gtcaagccgg cgctggttgc agaggtgcgc tacagcgagt ggactccgga
     2341 gggccggctg cgtcaatcaa gctggcgtgg gctgcggccg gacaagaaac ccagtgaggt
     2401 ggtgcgcgaa tgaattaaag aggagaatac tagatgcgag ccatttggac gggttcgatc
     2461 gccttcgggc tggtgaacgt gccggtcaag gtgtacagcg ctaccgcaga ccacgacatc
```

2521 aggttccacc aggtgcacgc caaggacaac ggacgcatcc ggtacaagcg cgtctgcgag
2581 gcgtgtggcg aggtggtcga ctaccgcgat cttgcccggg cctacgagtc cggcgacggc
2641 caaatggtgg cgatcaccga cgacgacatc gccagcttgc ctgaagaacg cagccgggag
2701 atcgaggtgt tggagttcgt ccccgccgcc gacgtggacc cgatgatgtt cgaccgcagc
2761 tactttttgg agcctgattc gaagtcgtcg aaatcgtatg tgctgctggc taagacactc
2821 gccgagaccg accggatggc gatcgtgcat ttcacgctgc gcaacaagac caggctggcg
2881 gcgttgcgcg tcaaggattt cggcaagcga gaggtgatga tggtgcacac gttgctgtgg
2941 cccgatgaga tccgcgaccc cgacttcccg gtgctggacc agaaggtgga gatcaaaccc
3001 gcggaactca agatggccgg ccaggtggtg gactcgatgg ccgacgactt caatccggac
3061 cgctaccacg acacctacca ggagcagtta caggagctga tcgacaccaa actcgaaggt
3121 gggcaggcat ttaccgccga ggaccaaccg aggttgctgg acgagcccga agacgtctcc
3181 gacctgctcg ccaagctgga ggccagcgtg aaggcgcgct cgaaggccaa ctcaaacgtc
3241 ccaacgcctc cgtgacgcat cctcacgata atatccgggt aggcgcaatc actttcgtct
3301 actccgttac aaagcgaggc tgggtatttc ccggcctttc tgttatccga aatccactga
3361 aagcacagcg gctggctgag gagataaata ataaacgagg ggctgtatgc acaaagcatc
3421 ttctgttgag ttaagaacga gtatcgagat ggcacatagc cttgctcaaa ttggaatcag
3481 gtttgtgcca ataccagtag aaacagacga agaatccatg ggtatggaca gcgcggaacc
3541 cctatttgtt tatttttcta aatacattca aatatgtatc cgctcatgag acaataaccc
3601 tgataaatgc ttcaataata ttgaaaaagg aagagtatga ttgaacaaga tggattgcac
3661 gcaggttctc cggccgcttg ggtggagagg ctattcggct atgactgggc acaacagaca
3721 atcggctgct ctgatgccgc cgtgttccgg ctgtcagcgc aggggcgccc ggttcttttt
3781 gtcaagaccg acctgtccgg tgccctgaat gaactgcagg acgaggcagc gcggctatcg
3841 tggctggcca cgacgggcgt tccttgcgca gctgtgctcg acgttgtcac tgaagcggga
3901 agggactggc tgctattggg cgaagtgccg gggcaggatc tcctgtcatc tcaccttgct
3961 cctgccgaga aagtatccat catggctgat gcaatgcggc ggctgcatac gcttgatccg
4021 gctacctgcc cattcgacca ccaagcgaaa catcgcatcg agcgagcacg tactcggatg
4081 gaagccggtc ttgtcgatca ggatgatctg gacgaagagc atcaggggct cgcgccagcc
4141 gaactgttcg ccaggctcaa ggcgcgcatg cccgacggcg aggatctcgt cgtgacccat
4201 ggcgatgcct gcttgccgaa tatcatggtg gaaaatggcc gcttttctgg attcatcgac
4261 tgtggccggc tgggtgtggc ggaccgctat caggacatag cgttggctac ccgtgatatt
4321 gctgaagagc ttggcggcga atgggctgac cgcttcctcg tgctttacgg tatcgccgct
4381 cccgattcgc agcgcatcgc cttctatcgc cttcttgacg agttcttctg aaaggaggtt
4441 ataaaaaatg aaaccagtaa cgttatacga tgtcgcagag tatgccggtg tctcttatca
4501 gaccgtttcc cgcgtggtga accaggccag ccacgtttct gcgaaaacgc gggaaaaagt
4561 ggaagcggcg atggcggagc tgaattacat tcccaaccgc gtggcacaac aactggcggg
4621 caaacagtcg ttgctgattg gcgttgccac ctccagtctg gccctgcacg cgccgtcgca
4681 aattgtcgcg gcgattaaat ctcgcgccga tcaactgggt gccagcgtgg tggtgtcgat
4741 ggtagaacga agcggcgtcg aagcctgtaa agcggcggtg cacaatcttc tcgcgcaacg
4801 cgtcagtggg ctgatcatta actatccgct ggatgaccag gatgccattg ctgtggaagc
4861 tgcctgcact aatgttccgg cgttatttct tgatgtctct gaccagacac ccatcaacag
4921 tattattttc tcccatgaag acggtacgcg actgggcgtg gagcatctgg tcgcattggg
4981 tcaccagcaa atcgcgctgt tagcgggccc attaagttct gtctcggcgc gtctgcgtct
5041 ggctggctgg cataaatatc tcactcgcaa tcaaattcag ccgatagcgg aacgggaagg

5101 cgactggagt gccatgtccg gttttcaaca aaccatgcaa atgctgaatg agggcatcgt
5161 tcccactgcg atgctggttg ccaacgatca gatggcgctg ggcgcaatgc gcgccattac
5221 cgagtccggg ctgcgcgttg gtgcggatat ctcggtagtg ggatacgacg ataccgaaga
5281 cagctcatgt tatatcccgc cgttaaccac catcaaacag gattttcgcc tgctggggca
5341 aaccagcgtg gaccgcttgc tgcaactctc tcagggccag gcggtgaagg gcaatcagct
5401 gttgcccgtc tcactggtga aaagaaaaac caccctggcg cccaatacgc aaaccgcctc
5461 tccccgcgcg ttggccgatt cattaatgca gctggcacga caggtttccc gactggaaag
5521 cgggcagtga taactgtcag accaagttta cgagctcgct tggactcctg ttgatagatc
5581 cagtaatgac ctcagaactc catctggatt tgttcagaac gctagggata acagggtaat
5641 tattgcgctg cagttttggg accattcaaa acagcatagc tctaaaacct cgtagactat
5701 ttttgtctaa aaaatttcgt aatcgcacta tttgtctcag ctagacttca gtcttgaaaa
5761 gcccctgtat tactgcattt attaagagta ttataccata tttttagtta ttaagaaata
5821 atcttcatct aaaatatact tcagtcacct cctagctgac tcaaatcaat gcgtgtttca
5881 taaagaccag tgatggattg atggataaga gtggcatcta aaacttcttt tgtagacgta
5941 tatcgtttac gatcaattgt tgtatcaaaa tatttaaaag cagcgggagc tccaagattc
6001 gtcaacgtaa ataaatgaat aatattttct gcttgttcac gtattggttt gtctctatgt
6061 ttgttatatg cactaagaac tttatctaaa ttggcatctg ctaaaataac acgcttagaa
6121 aattcactga tttgctcaat aatctcatct aaataatgct tatgctgctc cacaaacaat
6181 tgtttttgtt cgttatcttc tggactaccc ttcaactttt cataatgact agctaaatat
6241 aaaaaattca catatttgct tggcagagcc agctcatttc ctttttgtaa ttctccggca
6301 ctagccagca tccgtttacg accgttttct aactcaaaaa gactatattt aggtagttta
6361 atgattaagt cttttttaac ttccttatat cctttagctt ctaaaaagtc aatcggattt
6421 ttttcaaagg aacttctttc cataattgtg atccctagta actctttaac ggattttaac
6481 ttcttcgatt tcccttttc caccttagca accactagga ctgaataagc taccgttgga
6541 ctatcaaaac caccatattt ttttggatcc cagtcttttt tacgagcaat aagcttgtcc
6601 gaatttcttt ttggtaaaat tgactccttg gagaatccgc ctgtctgtac ttctgttttc
6661 ttgacaatat tgacttgggg catggacaat actttgcgca ctgtggcaaa atctcgccct
6721 ttatcccaga caatttctcc agtttcccca ttagtttcga ttagagggcg tttgcgaatc
6781 tctccatttg caagtgtaat ttctgttttg aagaagttca tgatattaga gtaaaagaaa
6841 tattttgcgg ttgctttgcc tatttcttgc tcagacttag caatcatttt acgaacatca
6901 taaactttat aatcaccata gacaaactcc gattcaagtt ttggatattt cttaatcaaa
6961 gcagttccaa cgacggcatt tagatacgca tcatgggcat gatggtaatt gttaatctca
7021 cgtactttat agaattggaa atcttttcgg aagtcagaaa ctaatttaga ttttaaggta
7081 atcactttaa cctctcgaat aagtttatca ttttcatcgt atttagtatt catgcgacta
7141 tccaaaattt gtgccacatg cttagtgatt tggcgagttt caaccaattg gcgtttgata
7201 aaaccagctt tatcaagttc actcaaacct ccacgttcag ctttcgttaa attatcaaac
7261 ttacgttgag tgattaactt ggcgtttaga agttgtctcc aatagttttt catctttttg
7321 actacttctt cacttggaac gttatccgat ttaccacgat ttttatcaga acgcgttaag
7381 accttattgt ctattgaatc gtctttaagg aaactttgtg gaacaatgtg atcgacatca
7441 taatcactta aacgattaat atctaattct tggtccacat acatgtctct tccattttgg
7501 agataataga gatagagctt ttcattttgc aattgagtat tttcaacagg atgctcttta
7561 agaatctgac ttcctaattc tttgatacct tcttcgattc gtttcatacg ctctcgcgaa
7621 tttttctggc ccttttgagt tgtctgattt tcacgtgcca tttcaataac gatattttct

7681 ggcttatgcc gccccattac tttgaccaat tcatcaacaa cttttacagt ctgtaaaata
7741 cctttttaa tagcagggct accagctaaa tttgcaatat gttcatgtaa actatcgcct
7801 tgtccagaca cttgtgcttt ttgaatgtct tctttaaatg tcaaactatc atcatggatc
7861 agctgcataa aattgcgatt ggcaaaacca tctgatttca aaaaatctaa tattgttttg
7921 ccagattgct tatccctaat accattaatc aattttcgag acaaacgtcc ccaaccagta
7981 taacggcgac gtttaagctg tttcatcacc ttatcatcaa agaggtgagc atatgtttta
8041 agtctttcct caatcatctc cctatcttca aataaggtca atgttaaaac aatatcctct
8101 aagatatctt cattttcttc attatccaaa aaatctttat ctttaataat ttttagcaaa
8161 tcatggtagg tacctaatga agcattaaat ctatcttcaa ctcctgaaat ttcaacacta
8221 tcaaaacatt ctattttttt gaaataatct tcttttaatt gcttaacggt tacttttcga
8281 tttgttttga agagtaaatc aacaatggct ttcttctgtt cacctgaaag aaatgctggt
8341 tttcgcattc cttcagtaac atatttgacc tttgtcaatt cgttataaac cgtaaaatac
8401 tcataaagca aactatgttt tggtagtact ttttcatttg gaagattttt atcaaagttt
8461 gtcatgcgtt caataaatga ttgagctgaa gcacctttat cgacaacttc ttcaaaattc
8521 catggggtaa ttgtttcttc agacttccga gtcatccatg caaaacgact attgccacgc
8581 gccaatggac caacataata aggaattcga aaagtcaaga tttttcaat cttctcacga
8641 ttgtctttta aaaatggata aaagtcttct tgtcttctca aaatagcatg cagctcaccc
8701 aagtgaattt gatggggaat agagccgttg tcaaaggtcc gttgcttgcg cagcaaatct
8761 tcacgattta gtttcaccaa taattcctca gtaccatcca tttttctaa aattggtttg
8821 ataaatttat aaaattcttc ttggctagct cccccatcaa tataacctgc atatccgttt
8881 tttgattgat caaaaaagat ttctttatac ttttctggaa gttgttgtcg aactaaagct
8941 tttaaaagag tcaagtcttg atgatgttca tcgtagcgtt taatcattga agctgatagg
9001 ggagccttag ttatttcagt atttactctt aggatatctg aaagtaaaat agcatctgat
9061 aaattcttag ctgccaaaaa caaatcagca tattgatctc caatttgcgc caataaatta
9121 tctaaatcat catcgtaagt atcttttgaa agctgtaatt tagcatcttc tgccaaatca
9181 aaatttgatt taaaattagg ggtcaaaccc aatgacaaag caatgagatt cccaaataag
9241 ccatttttct tctcaccggg gagctgagca atgagatttt ctaatcgtct tgatttactc
9301 aatcgtgcag aaagaatcgc tttagcatct actccacttg cgttaatagg gttttcttca
9361 aataattgat tgtaggtttg taccaactgg ataaatagtt tgtccacatc actattatca
9421 ggatttaaat ctccctcaat caaaaaatga ccacgaaact taatcatatg cgctaaggcc
9481 aaatagatta agcgcaaatc cgctttatca gtagaatcta ccaatttttt tcgcagatga
9541 tagatagttg gatatttctc atgataagca acttcatcta ctatatttcc aaaaatagga
9601 tgacgttcat gcttcttgtc ttcttccacc aaaaaagact cttcaagtcg atgaaagaaa
9661 ctatcatcta ctttcgccat ctcatttgaa aaaatctcct gtagataaca aatacgattc
9721 ttccgacgtg tataccttct acgagctgtc cgtttgagac gagtcgcttc cgctgtctct
9781 ccactgtcaa ataaaagagc ccctataaga ttttttga tactgtggcg gtctgtattt
9841 cccagaacct tgaacttttt agacggaacc ttatattcat cagtgatcac cgcccatccg
9901 acgctatttg tgccgatatc taagccttatt gagtatttct tatccatttt ttataacctc
9961 cttagagctc gaattcccaa aaaaacgggt atggagaaac agtagagagt tgcgataaaa
10021 agcgtcaggt aggatccgct aatcttatgg ataaaaatgc tatggcatag caaagtgtga
10081 cgccgtgcaa ataatcaatg tggactttc tgccgtgatt atagacactt ttgttacgcg
10141 tttttgtcat ggctttggtc ccgctttgtt acagaatgct tttaataagc ggggttaccg
10201 gtttggttag cgagaagagc cagtaaaaga cgcagtgacg gcaatgtctg atgcaatatg

10261 gacaattggt ttcttctctg aatggcggga gtatgaaaag tatggctgaa gcgcaaaatg
10321 atcccctgct gccgggatac tcgtttaatg cccatctggt ggcgggttta acgccgattg
10381 aggccaacgg ttatctcgat ttttttatcg accgaccgct gggaatgaaa ggttatattc
10441 tcaatctcac cattcgcggt cagggggtgg tgaaaaatca gggacgagaa tttgtttgcc
10501 gaccgggtga tattttgctg ttcccgccag gagagattca tcactacggt cgtcatccgg
10561 aggctcgcga atggtatcac cagtgggttt actttcgtcc gcgcgcctac tggcatgaat
10621 ggcttaactg gccgtcaata tttgccaata cggggttctt tcgcccggat gaagcgcacc
10681 agccgcattt cagcgacctg tttgggcaaa tcattaacgc cgggcaaggg gaagggcgct
10741 attcggagct gctggcgata aatctgcttg agcaattgtt actgcggcgc atggaagcga
10801 ttaacgagtc gctccatcca ccgatggata atcgggtacg cgaggcttgt cagtacatca
10861 gcgatcacct ggcagacagc aattttgata tcgccagcgt cgcacagcat gtttgcttgt
10921 cgccgtcgcg tctgtcacat cttttccgcc agcagttagg gattagcgtc ttaagctggc
10981 gcgaggacca acgtatcagc caggcgaagc tgcttttgag caccacccgg atgcctatcg
11041 ccaccgtcgg tcgcaatgtt ggttttgacg atcaactcta tttctcgcgg gtatttaaaa
11101 aatgcaccgg ggccagcccg agcgagttcc gtgccggttg tgaagaaaaa gtgaatgatg
11161 tagccgtcaa gttgtcataa taaatcgatg caggtggcac ttttcgggga aatgtggagg
11221 catcaaataa aacgaaaggc tcagtcgaaa gactgggcct ttcgttttat ctgttgtttg
11281 tcggtgaacg ctctcctgag taggacaaat ccgccgccct agacctaggg cgttcggctg
11341 cggcgagcgg tatcagctca ctcaaaggcg gtaatacggt tatccacaga atcaggggat
11401 aacgcaggaa agagcatgtg agcaaaaggc cagcaaaagg ccaggaaccg tggatatatt
11461 ccgcttcctc gctcactgac tcgctacgct cggtcgttcg actgcggcga gcggaaatgg
11521 cttacgaacg gggcggagat ttcctggaag atgccaggaa gatacttaac agggaagtga
11581 gagggccgcg gcaaagccgt ttttccatag gctccgcccc cctgacaagc atcacgaaat
11641 ctgacgctca aatcagtggt ggcgaaaccc gacaggacta taaagatacc aggcgtttcc
11701 ccctggcggc tccctcgtgc gctctcctgt tcctgccttt cggtttaccg gtgtcattcc
11761 gctgttatgg ccgcgtttgt ctcattccac gcctgacact cagttccggg taggcagttc
11821 gctccaagct ggactgtatg cacgaacccc ccgttcagtc cgaccgctgc gccttatccg
11881 gtaactatcg tcttgagtcc aacccggaaa gacatgcaaa agcaccactg gcagcagcca
11941 ctggtaattg atttagagga gttagtcttg aagtcatgcg ccggttaagg ctaaactgaa
12001 aggacaagtt ttggtgactg cgctcctcca agccagttac ctcggttcaa agagttggta
12061 gctcagagaa ccttcgaaaa accgccctgc aaggcggttt tttcgttttc agagcaagag
12121 attacgcgca gaccaaaacg atctcaagaa gatcatctta ttaataagga tctcaagaag
12181 atcctttgat cttttctacg gggtctgacg ctcagtggaa cgaaaactca cgttaaggga
12241 ttttggtcat gactagtgct tggattctca ccaataaaaa acgcccggcg gcaaccgatt
12301 tcaagttgat aacggactag ccttatttta acttgctatg ctgttttgaa tggttccaac
12361 aagattattt tataactttt ataacaaata atcaaggaga aattcaaaga aatttatcag
12421 ccgtgtcgcc cttaattgtg agcggataac aattacgagc ttcatgcaca gtgaaa

//

### Note S6. Complete sequence of the plasmid p-$P_{BAD}$-*cas9*/$P_{T5}$-Redγβα in Genbank format

LOCUS p-$P_{BAD}$-*cas9*/$P_{T5}$-Redγβα 11239 bp DNA circular SYN 24-NOV-2018
DEFINITION p-$P_{BAD}$-*cas9*/$P_{T5}$-Redγβα
ACCESSION p-$P_{BAD}$-*cas9*/$P_{T5}$-Redγβα
KEYWORDS
SOURCE
ORGANISM

FEATURES Location/Qualifiers
CDS 6..791
/label="Beta"
/note="single-stranded DNA binding recombinase in the λ Red system"
CDS 788..1468
/label="Exo"
/note="5' to 3' double-stranded DNA exonuclease in the λ Red system"
terminator 1469..1713
/label="lambda tL3 terminator"
promoter 1745..1849
/label="AmpR promoter"
CDS 1850..2644
/label=KanR
/note="aminoglycoside phosphotransferase from Tn5"
CDS 2661..3743
/label="lacI"
/note="lac repressor"
terminator 3776..3835
/label="lamda t0 terminator"
CDS complement(4054..8160)
/label="xCas9-3.7"
/note="generates RNA-guided double strand breaks in DNA"
promoter complement(8204..8369)
/label="araBAD promoter"
/note="promoter of the L-arabinose operon of E. coli"
CDS 8515..9393
/label="araC"
/note="L-arabinose regulatory protein"
terminator 9437..9483
/label="rrn B T1 terminator"
rep_origin complement(9815..10360)
/label="p15A ori"
/note="replication origin"

promoter 10690..10739

/label="T5 promoter"

CDS 10823..11239

/label="Gam"

/translation="inhibitor of the host RecBCD nuclease in the λ Red system"

ORIGIN

1 aacgaatgag tactgcactc gcaacgctgg ctgggaagct ggctgaacgt gtcggcatgg

61 attctgtcga cccacaggaa ctgatcacca ctcttcgcca gacggcattt aaaggtgatg

121 ccagcgatgc gcagttcatc gcattactga tcgttgccaa ccagtacggc cttaatccgt

181 ggacgaaaga aatttacgcc tttcctgata agcagaatgg catcgttccg gtggtgggcg

241 ttgatggctg gtcccgcatc atcaatgaaa accagcagtt tgatggcatg gactttgagc

301 aggacaatga atcctgtaca tgccggattt accgcaagga ccgtaatcat ccgatctgcg

361 ttaccgaatg gatggatgaa tgccgccgcg aaccattcaa aactcgcgaa ggcagagaaa

421 tcacggggcc gtggcagtcg catcccaaac ggatgttacg tcataaagcc atgattcagt

481 gtgcccgtct ggccttcgga tttgctggta tctatgacaa ggatgaagcc gagcgcattg

541 tcgaaaatac tgcatacact gcagaacgtc agccggaacg cgacatcact ccggttaacg

601 atgaaaccat gcaggagatt aacactctgc tgatcgccct ggataaaaca tgggatgacg

661 acttattgcc gctctgttcc cagatatttc gccgcgacat tcgtgcatcg tcagaactga

721 cacaggccga agcagtaaaa gctcttggat tcctgaaaca gaaagccgca gagcagaagg

781 tggcagcatg acaccggaca ttatcctgca gcgtaccggg atcgatgtga gagctgtcga

841 acagggggat gatgcgtggc acaaattacg gctcggcgtc atcaccgctt cagaagttca

901 caacgtgata gcaaaacccc gctccggaaa gaagtggcct gacatgaaaa tgtcctactt

961 ccacaccctg cttgctgagg tttgcaccgg tgtggctccg gaagttaacg ctaaagcact

1021 ggcctgggga aaacagtacg agaacgacgc cagaaccctg tttgaattca cttccggcgt

1081 gaatgttact gaatccccga tcatctatcg cgacgaaagt atgcgtaccg cctgctctcc

1141 cgatggttta tgcagtgacg gcaacggcct tgaactgaaa tgcccgttta cctcccggga

1201 tttcatgaag ttccggctcg gtggtttcga ggccataaag tcagcttaca tggcccaggt

1261 gcagtacagc atgtgggtga cgcgaaaaaa tgcctggtac tttgccaact atgacccgcg

1321 tatgaagcgt gaaggcctgc attatgtcgt gattgagcgg gatgaaaagt acatggcgag

1381 ttttgacgag atcgtgccgg agttcatcga aaaaatggac gaggcactgg ctgaaattgg

1441 ttttgtattt ggggagcaat ggcgatgacg catcctcacg ataatatccg ggtaggcgca

1501 atcactttcg tctactccgt tacaaagcga ggctgggtat ttcccggcct ttctgttatc

1561 cgaaatccac tgaaagcaca gcggctggct gaggagataa ataataaacg aggggctgta

1621 tgcacaaagc atcttctgtt gagttaagaa cgagtatcga gatggcacat agccttgctc

1681 aaattggaat caggtttgtg ccaataccag tagaaacaga cgaagaatcc atgggtatgg

1741 acagcgcgga acccctattt gtttattttt ctaaatacat tcaaatatgt atccgctcat

1801 gagacaataa ccctgataaa tgcttcaata atattgaaaa aggaagagta tgattgaaca

1861 agatggattg cacgcaggtt ctccggccgc ttgggtggag aggctattcg gctatgactg

1921 ggcacaacag acaatcggct gctctgatgc cgccgtgttc cggctgtcag cgcaggggcg

1981 cccggttctt tttgtcaaga ccgacctgtc cggtgccctg aatgaactgc aggacgaggc

2041 agcgcggcta tcgtggctgg ccacgacggg cgttccttgc gcagctgtgc tcgacgttgt

2101 cactgaagcg ggaagggact ggctgctatt gggcgaagtg ccggggcagg atctcctgtc

2161 atctcacctt gctcctgccg agaaagtatc catcatggct gatgcaatgc ggcggctgca

2221 tacgcttgat ccggctacct gcccattcga ccaccaagcg aaacatcgca tcgagcgagc
2281 acgtactcgg atggaagccg gtcttgtcga tcaggatgat ctggacgaag agcatcaggg
2341 gctcgcgcca gccgaactgt tcgccaggct caaggcgcgc atgcccgacg gcgaggatct
2401 cgtcgtgacc catggcgatg cctgcttgcc gaatatcatg gtggaaaatg gccgcttttc
2461 tggattcatc gactgtggcc ggctgggtgt ggcggaccgc tatcaggaca tagcgttggc
2521 tacccgtgat attgctgaag agcttggcgg cgaatgggct gaccgcttcc tcgtgcttta
2581 cggtatcgcc gctcccgatt cgcagcgcat cgccttctat cgccttcttg acgagttctt
2641 ctgaaaggag gttataaaaa atgaaaccag taacgttata cgatgtcgca gagtatgccg
2701 gtgtctctta tcagaccgtt tcccgcgtgg tgaaccaggc cagccacgtt tctgcgaaaa
2761 cgcgggaaaa agtggaagcg gcgatggcgg agctgaatta cattcccaac cgcgtggcac
2821 aacaactggc gggcaaacag tcgttgctga ttggcgttgc cacctccagt ctggccctgc
2881 acgcgccgtc gcaaattgtc gcggcgatta aatctcgcgc cgatcaactg ggtgccagcg
2941 tggtggtgtc gatggtagaa cgaagcggcg tcgaagcctg taaagcggcg gtgcacaatc
3001 ttctcgcgca acgcgtcagt gggctgatca ttaactatcc gctggatgac caggatgcca
3061 ttgctgtgga agctgcctgc actaatgttc cggcgttatt tcttgatgtc tctgaccaga
3121 cacccatcaa cagtattatt ttctcccatg aagacggtac gcgactgggc gtggagcatc
3181 tggtcgcatt gggtcaccag caaatcgcgc tgttagcggg cccattaagt tctgtctcgg
3241 cgcgtctgcg tctggctggc tggcataaat atctcactcg caatcaaatt cagccgatag
3301 cggaacggga aggcgactgg agtgccatgt ccggttttca acaaaccatg caaatgctga
3361 atgagggcat cgttcccact gcgatgctgg ttgccaacga tcagatggcg ctgggcgcaa
3421 tgcgcgccat taccgagtcc gggctgcgcg ttggtgcgga tatctcggta gtgggatacg
3481 acgataccga agacagctca tgttatatcc cgccgttaac caccatcaaa caggattttc
3541 gcctgctggg gcaaaccagc gtggaccgct tgctgcaact ctctcagggc caggcggtga
3601 agggcaatca gctgttgccc gtctcactgg tgaaaagaaa aaccaccctg gcgcccaata
3661 cgcaaaccgc ctctccccgc gcgttggccg attcattaat gcagctggca cgacaggttt
3721 cccgactgga aagcgggcag tgataactgt cagaccaagt ttacgagctc gcttggactc
3781 ctgttgatag atccagtaat gacctcagaa ctccatctgg atttgttcag aacgctaggg
3841 ataacagggt aattattgcg ctgcagtttt gggaccattc aaaacagcat agctctaaaa
3901 cctcgtagac tatttttgtc taaaaaattt cgtaatcgca ctatttgtct cagctagact
3961 tcagtcttga aaagcccctg tattactgca tttattaaga gtattatacc atatttttag
4021 ttattaagaa ataatcttca tctaaaatat acttcagtca cctcctagct gactcaaatc
4081 aatgcgtgtt tcataaagac cagtgatgga ttgatggata agagtggcat ctaaaacttc
4141 ttttgtagac gtatatcgtt tacgatcaat tgttgtatca aaatatttaa aagcagcggg
4201 agctccaaga ttcgtcaacg taaataaatg aataatattt tctgcttgtt cacgtattgg
4261 tttgtctcta tgtttgttat atgcactaag aactttatct aaattggcat ctgctaaaat
4321 aacacgctta gaaaattcac tgatttgctc aataatctca tctaaataat gcttatgctg
4381 ctccacaaac aattgttttt gttcgttatc ttctggacta cccttcaact tttcataatg
4441 actagctaaa tataaaaaat tcacatattt gcttggcaga gccagctcat ttcctttttg
4501 taacactccg gcactagcca gcatccgttt acgaccgttt tctaactcaa aaagactata
4561 tttaggtagt ttaatgatta agtctttttt aacttcctta tatcctttag cttctaaaaa
4621 gtcaatcgga ttttttcaa aggaacttct ttccataatt gtgatcccta gtaactcttt
4681 aacggatttt aacttcttcg atttcccttt ttccacctta gcaaccacta ggactgaata
4741 agctaccgtt ggactatcaa aaccaccata ttttttgga tcccagtctt ttttacgagc

4801 aataagcttg tccgaatttc tttttggtaa aattgactcc ttggagaatc cgcctgtctg
4861 tacttctgtt ttcttgacaa tattgacttg gggcatggac aatactttgc gcactgtggc
4921 aaaatctcgc cctttatccc agacaatttc tccagtttcc ccattagttt cgattagagg
4981 gcgtttgcga atctctccat ttgcaagtgt aatttctgtt ttgaagaagt tcatgatatt
5041 agagtaaaag aaatattttg cggttgcttt gcctatttct tgctcagact tagcaatcat
5101 tttacgaaca tcataaactt tataatcacc atagacaaac tccgattcaa gttttggata
5161 tttcttaatc aaagcagttc caacgacggc atttagatac gcatcatggg catgatggta
5221 attgttaatc tcacgtactt tatagaattg gaaatctttt cggaagtcag aaactaattt
5281 agattttaag gtaatcactt taacctctcg aataagttta tcattttcat cgtatttagt
5341 attcatgcga ctatccaaaa tttgtgccac atgcttagtg atttggcgag tttcaaccaa
5401 ttggcgtttg ataaaaccag ctttatcaag ttcactcaaa cctccacgtt cagctttcgt
5461 taaattatca aacttacgtt gagtgattaa cttggcgttt agaagttgtc tccaatagtt
5521 tttcatcttt ttgactactt cttcacttgg aacgttatcc gatttaccac gatttttatc
5581 agaacgcgtt aagaccttat tgtctattga atcgtcttta aggaaacttt gtggaacaat
5641 gtgatcgaca tcataatcac ttaaacgatt aatatctaat tcttggtcca catacatgtc
5701 tcttccattt tggagataat agagatagag cttttcattt tgcaattgag tattttcaac
5761 aggatgctct ttaagaatct gacttcctaa ttctttgata ccttcttcga ttcgtttcat
5821 acgctctcgc gaatttttct ggcccttttg agttgtctga ttttcacgtg ccatttcaat
5881 aacgatattt tctggcttat gccgccccat tactttgacc aattcatcaa caacttttac
5941 agtctgtaaa atacctttttt taatagcagg gctaccagct aaatttgcaa tatgttcatg
6001 taaactatcg ccttgtccag acacttgtgc tttttgaatg tcttctttaa atgtcaaact
6061 atcatcatgg atcagctgaa taaaattgcg attggcaaaa ccatctgatt tcaaaaaatc
6121 taatattgtt ttgccagatt gcttatccct aataccatta atcaattttc gagacaaacg
6181 tccccaacca gtataacggc gacgtttaag ctgtttcatc accttatcat caaagaggtg
6241 agcatatgtt ttaagtcttt cctcaatcat ctccctatct tcaaataagg tcaatgttaa
6301 aacaatatcc tctaagatat cttcattttc ttcattatcc aaaaaatctt tatctttaat
6361 aatttttagc aaatcatggt aggtacctaa tgaagcatta aatctatctt caactcctga
6421 aatttcaaca ctatcaaaac attctatttt tttgaaataa tcttctttta attgcttaac
6481 ggttactttt cgatttgttt tgaagagtaa atcaacaatg gctttcttct gatcacctga
6541 aagaaatgct ggttttcgca ttccttcagt aacatatttg acctttgtca attcgttata
6601 aaccgtaaaa tactcataaa gcaaactatg ttttggtagt actttttcat ttggaagatt
6661 tttatcaaag tttgtcatgc gttcaataaa tgattgagct gaagcacctt tatcgacaac
6721 tttttcaaaa ttccatgggg taattgtttc ttcagacttc cgagtcatcc atgcaaaacg
6781 actattgcca cgcgccaatg gaccaacata ataaggaatt cgaaaagtca agatttttttc
6841 aatcttctca cgattgtctt ttaaaaatgg ataaaagtct tcttgtcttc tcaaaatagc
6901 atgcagctca cccaagtgaa tttgatgggg aataatgccg ttgtcaaagg tccgttgctt
6961 gcgcagcaaa tcttcacgat ttagtttcac caataattcc tcagtaccat ccattttttc
7021 taaaattggt ttgataaatt tataaaattc ttcttggcta gctcccccat caatataacc
7081 tgcatatccg ttttttgatt gatcaaaaaa gatttcttta tacttttctg gaagttgttg
7141 tcgaactaaa gcttttaaaa gagtcaagtc ttgatgatgt tcatcgtaca gtttaatcat
7201 tgaagctgat aggggagcct tagttatttc agtatttact cttaggatat ctgaaagtaa
7261 aatagcatct gataaattct tagctgccaa aaacaaatca gcatattgat ctccaatttg
7321 cgccaataaa ttatctaaat catcatcgta agtatctttt gaaagctgta atttggtatc

7381 ttctgccaaa tcaaaatttg atttaaaatt aggggtcaaa cccaatgaca aagcaatgag
7441 attcccaaat aagccatttt tcttctcacc ggggagctga gcaatgagat tttctaatcg
7501 tcttgattta ctcaatcgtg cagaaagaat cgctttagca tctactccac ttgcgttaat
7561 agggttttct tcaaataatt gattgtaggt ttgtaccaac tggataaata gtttgtccac
7621 atcactatta tcaggattta aatctccctc aatcaaaaaa tgaccacgaa acttaatcat
7681 atgcgctaag gccaaataga ttaagcgcaa atccgcttta tcagtagaat ctaccaattt
7741 ttttcgcaga tgatagatag ttggatattt ctcatgataa gcaacttcat ctactatatt
7801 tccaaaaata ggatgacgtt catgcttctt gtcttcttcc accaaaaaag actcttcaag
7861 tcgatgaaag aaactatcat ctactttcgc catctcattt gaaaaaatct cctgtagata
7921 acaaatacga ttcttccgac gtgtatacct tctacgagct gtccgtttga gacgagtcgc
7981 ttccgctgtc tctccactgt caaataaaag agcccctata agattttttt tgatactgtg
8041 gcggtctgta tttcccagaa ccttgaactt tttagacgga accttatatt catcagtgat
8101 caccgcccat ccgacgctat ttgtgccgat atctaagcct attgagtatt tcttatccat
8161 tttttataac ctccttagag ctcgaattcc caaaaaaacg ggtatggaga aacagtagag
8221 agttgcgata aaaagcgtca ggtaggatcc gctaatctta tggataaaaa tgctatggca
8281 tagcaaagtg tgacgccgtg caaataatca atgtggactt ttctgccgtg attatagaca
8341 cttttgttac gcgtttttgt catggctttg gtcccgcttt gttacagaat gcttttaata
8401 agcggggtta ccggtttggt tagcgagaag agccagtaaa agacgcagtg acggcaatgt
8461 ctgatgcaat atggacaatt ggtttcttct ctgaatggcg ggagtatgaa aagtatggct
8521 gaagcgcaaa atgatcccct gctgccggga tactcgttta atgcccatct ggtggcgggt
8581 ttaacgccga ttgaggccaa cggttatctc gatttttta tcgaccgacc gctgggaatg
8641 aaaggttata ttctcaatct caccattcgc ggtcaggggg tggtgaaaaa tcagggacga
8701 gaatttgttt gccgaccggg tgatattttg ctgttcccgc caggagagat tcatcactac
8761 ggtcgtcatc cggaggctcg cgaatggtat caccagtggg tttactttcg tccgcgcgcc
8821 tactggcatg aatggcttaa ctggccgtca atatttgcca atacggggtt ctttcgcccg
8881 gatgaagcgc accagccgca tttcagcgac ctgtttgggc aaatcattaa cgccgggcaa
8941 ggggaagggc gctattcgga gctgctggcg ataaatctgc ttgagcaatt gttactgcgg
9001 cgcatggaag cgattaacga gtcgctccat ccaccgatgg ataatcgggt acgcgaggct
9061 tgtcagtaca tcagcgatca cctggcagac agcaattttg atatcgccag cgtcgcacag
9121 catgtttgct tgtcgccgtc gcgtctgtca catcttttcc gccagcagtt agggattagc
9181 gtcttaagct ggcgcgagga ccaacgtatc agccaggcga agctgctttt gagcaccacc
9241 cggatgccta tcgccaccgt cggtcgcaat gttggttttg acgatcaact ctatttctcg
9301 cgggtattta aaaaatgcac cggggccagc ccgagcgagt tccgtgccgg ttgtgaagaa
9361 aaagtgaatg atgtagccgt caagttgtca taataaatcg atgcaggtgg cacttttcgg
9421 ggaaatgtgg aggcatcaaa taaaacgaaa ggctcagtcg aaagactggg cctttcgttt
9481 tatctgttgt ttgtcggtga acgctctcct gagtaggaca aatccgccgc cctagaccta
9541 gggcgttcgg ctgcggcgag cggtatcagc tcactcaaag gcggtaatac ggttatccac
9601 agaatcaggg gataacgcag gaaagagcat gtgagcaaaa ggccagcaaa aggccaggaa
9661 ccgtggatat attccgcttc ctcgctcact gactcgctac gctcggtcgt tcgactgcgg
9721 cgagcggaaa tggcttacga acggggcgga gatttcctgg aagatgccag gaagatactt
9781 aacagggaag tgagagggcc gcggcaaagc cgtttttcca taggctccgc ccccctgaca
9841 agcatcacga aatctgacgc tcaaatcagt ggtggcgaaa cccgacagga ctataaagat
9901 accaggcgtt tccccctggc ggctccctcg tgcgctctcc tgttcctgcc tttcggttta

9961 ccggtgtcat tccgctgtta tggccgcgtt tgtctcattc cacgcctgac actcagttcc
10021 gggtaggcag ttcgctccaa gctggactgt atgcacgaac cccccgttca gtccgaccgc
10081 tgcgccttat ccggtaacta tcgtcttgag tccaacccgg aaagacatgc aaaagcacca
10141 ctggcagcag ccactggtaa ttgatttaga ggagttagtc ttgaagtcat gcgccggtta
10201 aggctaaact gaaaggacaa gttttggtga ctgcgctcct ccaagccagt tacctcggtt
10261 caaagagttg gtagctcaga gaaccttcga aaaaccgccc tgcaaggcgg ttttttcgtt
10321 ttcagagcaa gagattacgc gcagaccaaa acgatctcaa gaagatcatc ttattaataa
10381 ggatctcaag aagatccttt gatcttttct acggggtctg acgctcagtg gaacgaaaac
10441 tcacgttaag ggattttggt catgactagt gcttggattc tcaccaataa aaaacgcccg
10501 gcggcaaccg atttcaagtt gataacggac tagccttatt ttaacttgct atgctgtttt
10561 gaatggttcc aacaagatta ttttataact tttataacaa ataatcaagg agaaattcaa
10621 agaaatttat cagccgtgtc gcccttaatt gtgagcggat aacaattacg agcttcatgc
10681 acagtgaaat catgaaaaat ttatttgctt tgtgagcgga taacaattat aatatgtgga
10741 attgtgagcg ctcacaattc cacaacggtt tccctctaga aataattttg tttaactttt
10801 cgagacctta ggaggtaaac atatggatat taatactgaa actgagatca agcaaaagca
10861 ttcactaacc ccctttcctg ttttcctaat cagcccggca tttcgcgggc gatattttca
10921 cagctatttc aggagttcag ccatgaacgc ttattacatt caggatcgtc ttgaggctca
10981 gagctgggcg cgtcactacc agcagctcgc ccgtgaagag aaagaggcag aactggcaga
11041 cgacatggaa aaaggcctgc cccagcacct gtttgaatcg ctatgcatcg atcatttgca
11101 acgccacggg gccagcaaaa aatccattac ccgtgcgttt gatgacgatg ttgagtttca
11161 ggagcgcatg gcagaacaca tccggtacat ggttgaaacc attgctcacc accaggttga
11221 tattgattca gaggtataa
//
